# The microbial landscape: soil microbiome properties predict plant species distributions

**DOI:** 10.1101/2025.10.14.682295

**Authors:** Daniel Revillini, Caitlin C. Mothes, Brianna K. Almeida, Katherine T. Charton, Stephanie Koontz, Aaron S. David, Michelle E. Afkhami, Christopher A. Searcy

## Abstract

Plant species distributions are shaped by interactions with the abiotic and biotic environment. Despite the known importance of soil microbiomes in shaping plant diversity and function, no study has explicitly determined the ability of the soil microbiome to predict plant species distributions at large scales. We employed paired above- and belowground surveys of plant occurrence and soil microbial taxa and functions across habitat patches (n = 676), applying machine-learning-based distribution modeling to identify the relative influence of the soil microbiome and environmental attributes in predicting plant species (n = 50) distributions across the landscape. We discovered that while abiotic gradients of known importance in this ecosystem were the strongest predictors of many plant species’ distributions, microbial predictors could have similar or greater influence. Microbiome predictors were collectively more important than abiotic environmental variables for 38% of plant species in this study and explained >70% of the predicted distribution for one species. We identified four microbiome attributes of landscape-scale importance for predicting plant species distributions including prokaryotic richness, fungal richness, the abundance of fungal pathogens, and the abundance of prokaryotic phosphate transport genes in soil. Our findings reveal a previously underappreciated role of the soil microbiome in shaping plant species distributions at a landscape scale, with implications for plant community structure in the context of both ecosystem restoration and future global change.

## Introduction

The known importance of the soil microbiome to community-, landscape- and ecosystem-level ecology has greatly expanded in recent years (Schnitzer *et al*., 2011; Delgado-Baquerizo *et al*., 2016; Trivedi *et al*., 2016; Yang *et al*., 2018; De Vries *et al*., 2020), and it is becoming apparent that to understand the ways plants will respond to global change we must consider the role of this ‘unseen majority’ belowground (van der Heijden *et al*., 2008; Anthony *et al*., 2023). For example, a novel analysis of microbial effects on plant demography recently revealed a critical role of plant species filtering by microbial mutualists that ultimately weakens the latitudinal diversity gradient on islands (Delavaux *et al*., 2024). In forests, soil microbial mediation of plant stress appears to allow the shift of tree species into novel ranges as they experience climate change, where microbial communities adapted to certain stress conditions can lead to increased survival of tree seedlings experiencing those new conditions (Afkhami, 2023; Allsup *et al*., 2023). Additionally, it is largely understood that microbial pathogens, typically fungi, in soil can structure the aboveground community through various mechanisms (Dobson & Crawley, 1994; Mills & Bever, 1998; Bever, 2002), including disease outbreaks or negative plant-fungal feedbacks (Callaway *et al*., 2004; Mangan *et al*., 2010; Bagchi *et al*., 2014). These studies highlight the important role of the microbiome in shaping plant species performance and community dynamics, but there remains a critical gap in describing the explicit potential of the soil microbiome to shape plant species distributions.

Species distributions are the product of tolerances to abiotic conditions and the outcomes of biotic interactions. With regard to biotic interactions shaping distributions, a basic principle suggests that a species’ niche can either be contracted through antagonistic (e.g., competition, predation, pathogenesis) interactions or expanded via beneficial or mutualistic interactions (Afkhami *et al*., 2014; Peay, 2016; Blonder, 2018). Given the diversity of well-documented roles of soil microbes in both positive and negative interactions with plants (e.g., fungal pathogens, archaeal and bacterial plant-growth promoters, mutualistic mycorrhizal fungi; (Westover *et al*., 2001; Rillig, 2004; Lugtenberg & Kamilova, 2009)), it begs the question, might plant distributions also be driven by these ‘hidden players’ in the soil? The microbiome can directly and indirectly (via mediation) alter plant establishment (e.g., germination), performance (e.g., productivity), fitness (e.g., survival), and reproduction (e.g., flowering or seed production), shifting the realized niche and ultimately affecting plant community diversity and ecosystem functions (Delgado-Baquerizo *et al*., 2016, 2020; Kandlikar *et al*., 2019; Wagg *et al*., 2019). Additionally, these microbial effects are shown to be particularly important for plants experiencing environmental change or stress (Liu *et al*., 2020; Trivedi *et al*., 2020), which suggests an increasingly important role of the soil microbiome in plant health, as anthropogenic global change impacts will continue to escalate for decades to come. Finally, it is important to recognize that while the outcomes of intricately coevolved relationships among plants and pathogens or mutualists (*e.g.*, mycorrhizal fungi) are generally well-understood (Kiers & Denison, 2008; Afkhami *et al*., 2020; Taylor *et al*., 2020), plants roots maintain many complex and often more diffuse interactions with the hyperdiverse and multifunctional soil microbiome, with unknown effects on aboveground species distributions. Here, we consider a bottom-up perspective regarding the soil microbiome role in plant species distributions to identify whether (and specifically which attributes of) the soil microbiome may play a previously unidentified, strong, and selective role in structuring aboveground species occurrences across an entire landscape.

In this study, we employed a combination of landscape-scale, paired plant and soil community surveys (n = 676), including determination of plant identity, targeted amplicon sequencing (16S, ITS, SSU) for microbial community analysis and functional annotation, and machine-learning based species distribution modeling (SDM) in order to determine if and when microbial biodiversity and functions can predict plant species distributions. We hypothesized that: 1) abiotic variables of known importance in our study system such as fire regime would be influential in shaping the distribution of many plant species, but also that 2) soil microbial diversity and critical nutrient cycling functions (e.g., nitrogen fixation or phosphate cycling) could exert strong effects on plant species distributions. Our study reveals that while key abiotic gradients such as time since last fire have the greatest influence on plant habitat occupancy, the soil microbiome also has strong potential to shape plant distributions at the landscape scale. We identify four microbial metrics that were consistently important in predicting species distributions, which presents a promising potential strategy to target and manage key soil microbial factors that could be critical for plant communities and assist ecosystem restoration that aims to maintain or augment aboveground biodiversity through the management of microbiome properties.

This is the first study to map above- and belowground communities at this scale to better understand the potential effects of the soil microbiome in shaping plant distributions. By integrating microbial taxonomic and functional data with plant occurrence across a heterogeneous landscape, our approach provides a framework for assessing how belowground communities contribute to aboveground biodiversity patterns. Understanding these relationships is especially important as global change progresses, factors such as atmospheric nitrogen deposition, severe drought, and increasingly frequent extreme weather events are expected to alter soil conditions (De Vries *et al*., 2012; Zhang *et al*., 2018; Jansson & Hofmockel, 2020; Qu *et al*., 2024), shifting the composition of key microbial taxa (e.g., altering richness) and microbiome functions (e.g., declines in nutrient cycling) that can significantly affect plant distributions (García-Palacios *et al*., 2012; Sayer *et al*., 2017; Rudgers *et al*., 2025), especially those that are endemic or endangered. Therefore, beyond international strategies to mitigate global change, we advocate for further research to understand the role of global changes on soil microbiome properties and subsequent feedbacks on plant communities.

## Methods

### Study system

The Florida Scrub ecosystem has the highest rate of endemism in the southeastern United States and hosts many threatened plant species (Menges & Kohfeldt, 1995; Menges *et al*., 2008). This ecosystem exhibits a range of habitat types from open sand gaps and shrublands to mixed conifer flatwoods within a relatively small area. Many of the endangered, rare, and endemic plants in this ecosystem are found in the rosemary scrub habitat, where they occur in open sand gaps, defined as areas with open canopies between shrubs such as the dominant, allelopathic Florida rosemary (*Ceratiola ericoides* Michaux) as well as oaks, ericads, and palmettos (Menges *et al*., 2008). Soil in the rosemary scrub is categorized as excessively well-drained Entisols, mostly comprised of the USDA-NRCS soil series Archbold and St. Lucie (Navarra *et al*., 2011). These white quartz sands are acidic (∼4 - 5.1 pH) and low in clay, organic matter, and nutrients (Abrahamson *et al*., 1984; Dean, 2015). Fire and elevation gradients are the main abiotic drivers of the patchy structure of this habitat type (Fischer *et al*., 1994; Menges & Kohfeldt, 1995; Dee & Menges, 2014). The habitat is managed using prescribed fire every 15 - 60 yr, and Archbold maintains a detailed database of fire history at the 5 x 5 m grid scale (Menges *et al*., 2017). The elevation gradients in this habitat have been shown to influence both plant and microbial communities, largely through the change in relative elevation to the water table, a proxy for soil moisture (Menges & Hawkes, 1998; Quintana-Ascencio *et al*., 2018; Hernandez *et al*., 2021). Recent studies have also demonstrated an important role of soil microbiome interactions influencing both plant productivity and fitness (David *et al*., 2018; Revillini *et al*., 2022, 2023).

### Soil sampling and plant surveys

Soil samples were collected from 676 open sand gaps (mean area = 8.3 m^2^, sd = 8.8 m^2^) between Florida rosemary shrubs from 28 balds (larger rosemary scrub units) across the Florida rosemary scrub landscape at Archbold Biological Station (Venus, FL, USA) using sterile technique for microbiome analyses (Figure 1). Specifically, three samples were collected from each gap (middle of the gap and halfway to the gap edge in opposite directions) and homogenized for a representative sample between 7-14 May, 2018. This sampling method specifically minimizes the influence of allelopathic Florida rosemary by staying away from gap edges, and represents the soil microbiome that would be experienced by germinating seeds in this ecosystem (in open sand gaps). Presence/absence for all plant species in this community was determined at the same 676 sites during the following growing season (September-December 2018). In total, 76 plant species were identified in our survey (Supplemental Figure 1). Lichens were included in the plant community due to their ecological relevance, consistent co-occurrence with vascular plants in this system, and their large amount of ground cover (Menges & Kohfeldt, 1995). The selected sites represent a comprehensive sampling across the two most important abiotic gradients influencing plant communities in this habitat: time since the last fire (TSF; 6 - 1427 months) and elevation above the water table (0.32 – 2.73 m).

**Figure 1.**
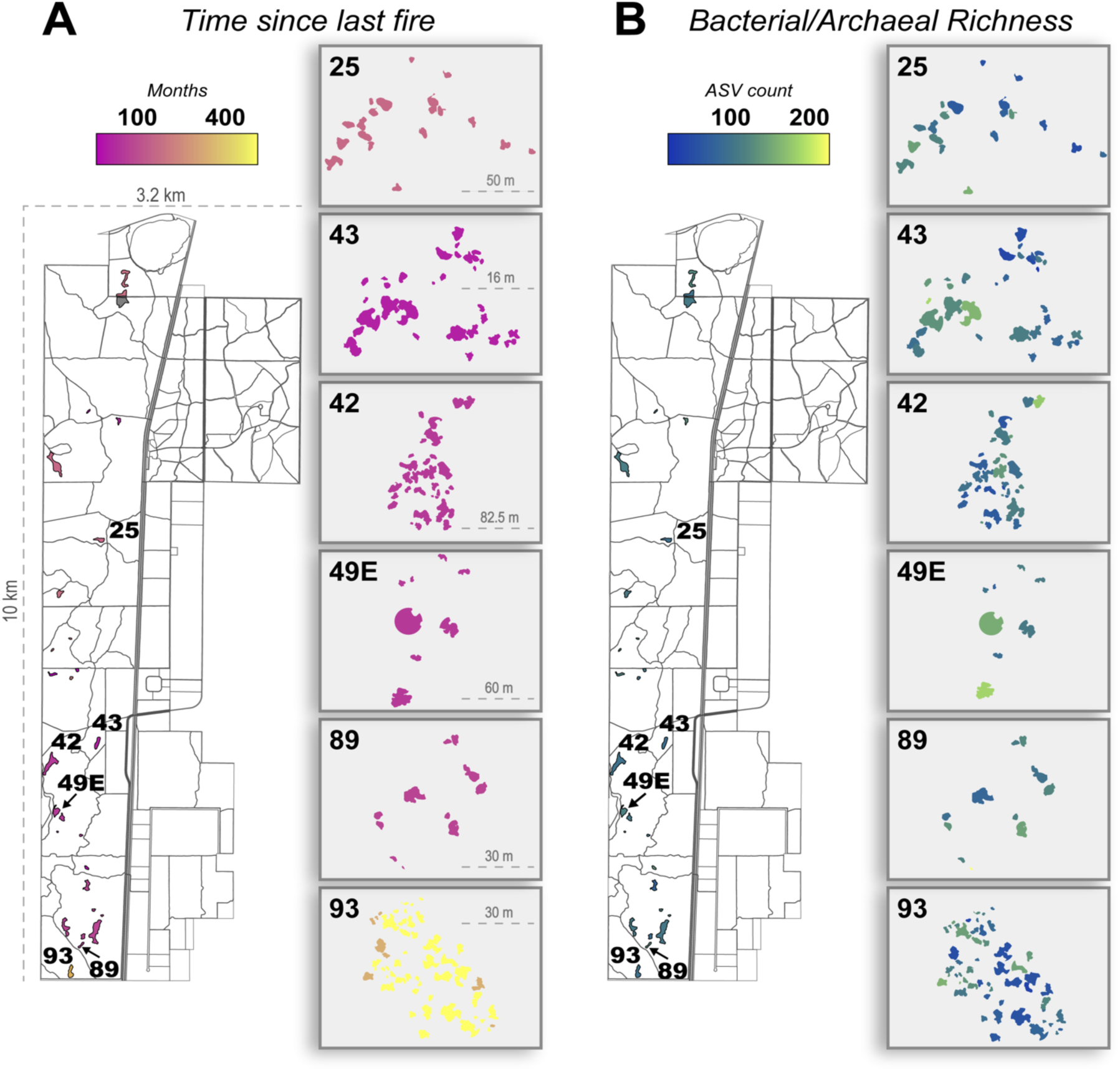
Maps of the **A)** abiotic predictor variable, time since last fire, and **B)** soil microbial predictor variable, taxonomic richness of soil prokaryotes, that each contributed most to explaining plant species distributions across the Florida rosemary scrub landscape at Archbold Biological Station (Venus, Florida, USA). In the full landscape maps, all rosemary scrub sites (balds) are colored by their average values and are labeled for associated blow-ups. Blow-ups are shown for six of the 28 balds in order to illustrate variation between the open sand gaps within balds. Most of variation in TSF was between balds, while most of the variation in prokaryotic richness was between open sand gaps within balds. Scale is provided for the landscape map and for each blow-up.

### Molecular analyses and bioinformatics

DNA was extracted from homogenized soil samples from each open sand gap (n = 676) using the DNeasy PowerSoil Pro QIAcube HT Kit (Qiagen, Carlsbad, CA, USA) with an adapted protocol without QIAcube (see Supporting *Detailed Methods* section for full description; and Revillini *et al*., 2022). DNA was quantified with a Qubit 4 fluorometer (Qiagen, Carlsbad, CA, USA) and normalized to 5 ng/μL. Libraries were prepared for sequencing using a two-step dual indexing protocol (Gohl *et al*., 2016). PCR was targeted for archaeal/bacterial (16S), broad fungal (ITS2), and arbuscular mycorrhizal fungi (SSU) ribosomal DNA using primer pairs 515F-806R, ITS7o-ITS4, and WANDA-AML2, respectively. Index and Illumina flowcell sequences were added in second-step PCR. All targeted amplicon products were pooled in equimolar quantities and sent to the Duke University Microbiome Core Facility (Durham, NC, USA). Libraries were sequenced on a MiSeq Desktop Sequencer (v3, 300 bp paired end; Illumina, Inc., San Diego, CA, USA).

Paired-end molecular sequence data was processed using QIIME2 v2021.4 (Bolyen *et al*., 2019). Briefly, denoising was performed with the DADA2 algorithm (Callahan *et al*., 2016). Naive Bayes classifiers were constructed using the Greengenes database v13.8 (99% similarity; *i.e.*, amplicon sequence variant (ASV)) and the UNITE database v7.2 (97% similarity; *i.e.*, operational taxonomic unit (OTU)) for prokaryotic and fungal amplicons, respectively. Taxa were classified using the sklearn algorithm. Multiple sequence alignments were performed using mafft v7 (Katoh & Standley, 2013), an unrooted tree was created using FastTree2, and the midpoint root method was used to create a rooted tree for weighted UniFrac analyses of prokayotes. Taxa not present in at least two samples were filtered out, and diversity metrics and dissimilarity matrices were calculated using the QIIME2 commands *diversity core-metrics-phylogenetic* (sampling depth = 6,500) and *diversity core-metrics* (sampling depth = 9,000) for archaea/bacteria and fungi, respectively. After quality control of sequencing data, 660 sampling sites were retained for further analysis. Microbiome data was read into R v4.4 (R Core Team, 2020) using the qiime2R package v0.99.6 (https://github.com/jbisanz/qiime2R).

### Microbiome metrics to predict plant species distributions

We calculated alpha-diversity metrics for each targeted microbial group: prokaryotic (16S), broad fungal (ITS), and AM fungal (SSU) to be used in species distribution models. Fungal functional classification was performed with ITS sequence data. The relative abundance of potential fungal pathogens, saprotrophs, and symbionts were determined using the FUNGuild database (Nguyen *et al*., 2016), and only using taxa whose trophic guild assignment was classified as “highly probable” or “probable”. Dominant taxa from each fungal trophic mode were determined based on greater than average relative abundance (read count > mean per trophic mode) and high occurrence across all sites (>50%), similar to determination of the “core microbiome” (Neu *et al*., 2021). Functional potential of bacterial/archaeal communities was determined by inferencing gene content from taxonomic abundances using an ASV approach with the PICRUSt2 algorithm (Douglas *et al*., 2020). All ASVs used for functional metagenomic prediction were below the recommended NSTI < 2.0 cutoff. For the species distribution models, we targeted analyses on genes associated with important carbon (C), nitrogen (N), and phosphorus (P)-cycling functions including individual genes (*e.g*., *nifQ* or *amoA*), or sums of functional gene sets/types representing functional pathways: N transport (*nrtP*, *narK*, *nasA*), nitrite reduction (*nirB, nirD, nirK*), phosphonate (organic P) cleavage and transport (*phnC, phnD, phnE, phnJ*), inorganic glycerol phosphate uptake and transport (*ugpA, ugpC, ugpQ*) and glucosidase degradation (*bglB* and *malZ*).

### Species distribution models

Boosted regression tree (BRT) models were generated for each of the plant species identified in ≥10 open sand gaps during the plant community surveys (n = 50). BRTs combine multiple simple regression tree models through an adaptive boosting process, creating a powerful ensemble approach that outperforms traditional single-model statistical methods (Elith *et al*., 2008) and have proved to be one of the best performing methods for species distribution models (Valavi *et al*., 2022). Our study employs a commonly used BRT implementation that utilizes stochastic gradient boosting methodology (Friedman, 2002). The *gbm.step()* function from the ‘dismo’ R package was used (Hijmans *et al*., 2024), with a Bernoulli distribution and a minimum of 1000 tree iterations per plant species. Sixteen variables were considered as potential drivers of plant species distributions: two abiotic gradients and 14 characterizing the soil microbiome. The two abiotic variables (elevation above the water table and TSF) are gradients with well-documented and important effects on plant community composition in this ecosystem (Dee & Menges, 2014; Hernandez *et al*., 2021; Revillini *et al*., 2022; Quintana-Ascencio *et al*., 2018). We included the two abiotic gradients in models to compare the influence of untested microbiome properties with the most well-known contributors to plant community assembly and structure in this ecosystem. We compiled the 14 microbial metrics from all soil samples (n = 660) to use as microbiome predictors of plant species distributions. These included observed taxonomic richness for three important soil microbial groups: prokaryotes (16S), all fungi (ITS), and arbuscular mycorrhizal fungi (SSU); the relative abundance of five prokaryotic N and P cycling genes/gene pathways: N-transport, nitrite reduction, orthophosphate uptake, P mineralization, and hemicellulose degradation (see above for details); and the relative abundance of taxa representing six fungal trophic modes: pathogens, saprotrophs, symbionts, pathogen-saprotrophs, saprotroph-symbionts, and pathogen-saprotroph-symbionts. The direction of each variable’s influence on probability of species occurrence was visualized with partial dependence plots from the *plot.gbm* function in R. Based on these plots, plant species were sorted into those that are most abundant at low, intermediate, or high values along each of the abiotic and biotic variables, with particular attention paid to variables that contributed more to each species’ probability of occurrence than expected by chance (*i.e*., variable importance > 1/16).

### Data analysis

Variation in predictor variables was determined using ANOVA at two spatial scales (bald and sand gaps within bald to determine whether predictor variance was greater within or between the largest sampling units. Generalized linear models were used to determine the relationships between the two abiotic gradients and 14 microbial properties and landscape-level plant richness of all plant species identified in surveys (n = 76), and the subset of 21 plants species with informative species distribution models (test area under curve (AUC) > 0.7 based on 10-fold cross validation) from the BRT analyses. Best-supported models were identified with backwards stepwise regression using the *step* function and partial F-test based on model cutoff at P < 0.05. The *Anova* function from the “car” package was used to summarize model variable significance using type III sums of squares (“type = III”). All analyses were performed in R version 4.4.1.

## Results

### Relationships between plant community diversity, time since last fire, and the soil microbiome

Microbiome predictors were more variable within balds, while abiotic predictors were more variable between balds (microbiome predictors: 80-84% of total variance within balds; abiotic predictors: 81-97% of total variance between balds; Figure 1). Site-level plant species richness was primarily associated with three soil microbiome predictors and TSF (Supplemental Figure 2 & Supplemental Figure 3). Generalized linear models revealed that estimated prokaryotic orthophosphate (Pi) uptake-and-transport genes abundance was the strongest predictor for all 76 plant species surveyed across the landscape, as well as the subset of 21 plant species with well-predicted distributions (F_1,642_= 14.7, *P* < 0.001; F_1,642_ = 15.5, *P* < 0.001, respectively). In general, plant richness was high in sites with low abundance of *ugp* genes. TSF was the second most important variable for both total plant richness and richness of the 21 species with well-predicted distributions (F_1,642_= 13.1, *P* < 0.001; F_1,642_ = 14.9, *P* < 0.001, respectively). A negative relationship between TSF and both total and well-predicted plant richness was observed (Supplemental Figure 2). Finally, fungal richness and the estimated abundance of prokaryotic nitrite reductase genes significantly explained the observed richness for both the total plant community (F_1,642_ = 11.18, *P* < 0.001; F_1,642_= 10.3, *P* = 0.001, respectively) and the 21 species that had well predicted distributions (F_1,642_ = 10.83, *P* = 0.001; F_1,642_= 8.04, *P* = 0.005, respectively), with plant richness increasing with both fungal richness and prokaryotic nitrite reductase genes abundance (Supplemental Figure 2).

### Abiotic factors and plant species distributions

To gain deeper insight into the role of environmental gradients and soil microbiome properties for individual plant species distributions, we conducted a detailed evaluation of the subset of plant species for which collected data allowed us to generate high quality SDMs. Of the 50 plant SDMs constructed, 21 species (42%) were well-predicted by the abiotic and microbial factors considered in our models (test AUC > 0.70; Table 1). Of the 21 informative models, seven species were forbs, six species were ground lichens, four were subshrubs, three were grasses, and one was a lycopod. TSF was the most important predictor for 18 of the 21 informative SDMs (Figure 2, Supplemental Figure 3) and the contribution was greater than expected by chance for all 21 informative SDMs (Supplemental Figure 4). Three lichen (*Cladonia evansii*, *Cladonia perforata* and *Cladonia prostrata*) and one lycopod (*Selaginella arenicola*) species were those most strongly influenced by TSF (78%, 52% 49% and 42%, respectively; Figure 2). Directionality of the TSF effect (from partial dependence plots) was well defined for 20 out of 21 species (Supplemental Figure 5). We observed that plant functional groups responded differently to TSF, with forbs and grasses favored in the first 12-13 years post-fire and lichens favored when the TSF interval exceeded ∼30 years. There is an intermediate TSF period during which the functional assemblage is more mixed. It is also notable that almost all of the endangered/endemic plants with well resolved models are more common shortly after fire, with the notable exception of *Cladonia perforata*, one of only two federally endangered lichens (Supplemental Figure 5). Relative elevation to the water table was the second strongest predictor across the 21 informative SDMs. Approximately 71% of the dependence curve peaks were well-defined for relative elevation to the water table (15 of 21), with roughly equal numbers of species specializing on each portion of this gradient (low, middle, and high), but with less clear separation between functional groups than for TSF. Relative elevation had the greatest variable importance for two plant species: *Clinipodium asheii* (35%) and *Andropogon floridanus* (34%) (Figure 2), both of which specialize on open sand gaps closest to the water table (<1.4 m).

**Figure 2.**
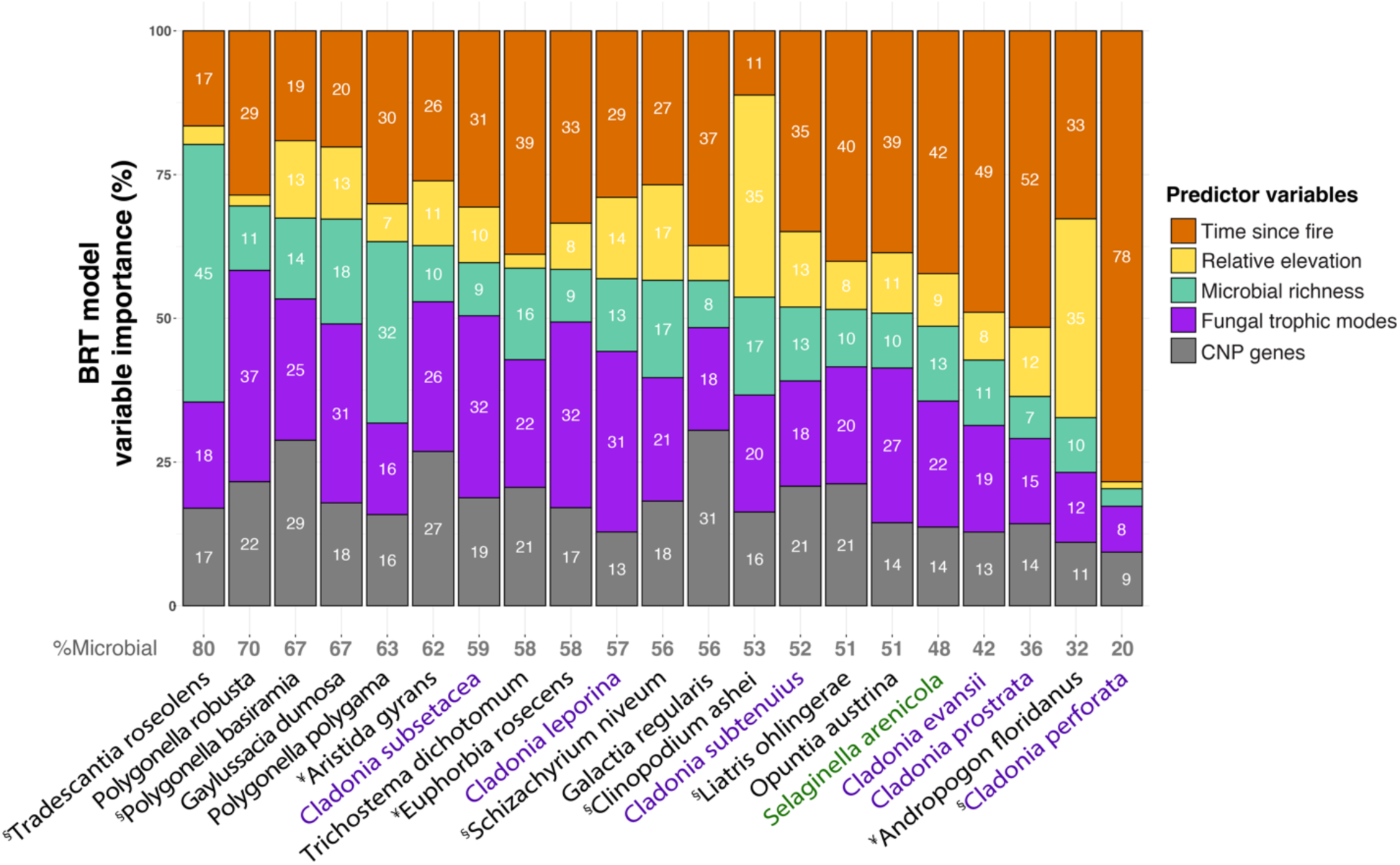
Predictor importance for all plant SDMs with test AUC > 0.70, including abiotic variables, time since last fire and relative elevation to water table, as well as categories of microbial predictors: microbial richness indicates 16S, ITS, and SSU observed taxonomic richness; fungal trophic modes indicate abundance of six classified fungal trophic modes; CNP genes indicates abundance of prokaryotic functional genes associated with carbon, nitrogen and phosphorus cycling. Total percent of each species’ distribution explained by microbial predictors is in gray (bottom), and plant species are arranged in descending order of this metric (%Microbial) along the x-axis. For 38% of plant species (eight of 21), a microbial predictor category explained the distribution more than either abiotic variable. Lichens are in purple and the lycopod (*Selaginella arenicola*) is in green. ¥ = plant species endemic to Florida or southeastern USA. § = listed as a threatened or endangered plant species.

**Table 1.**
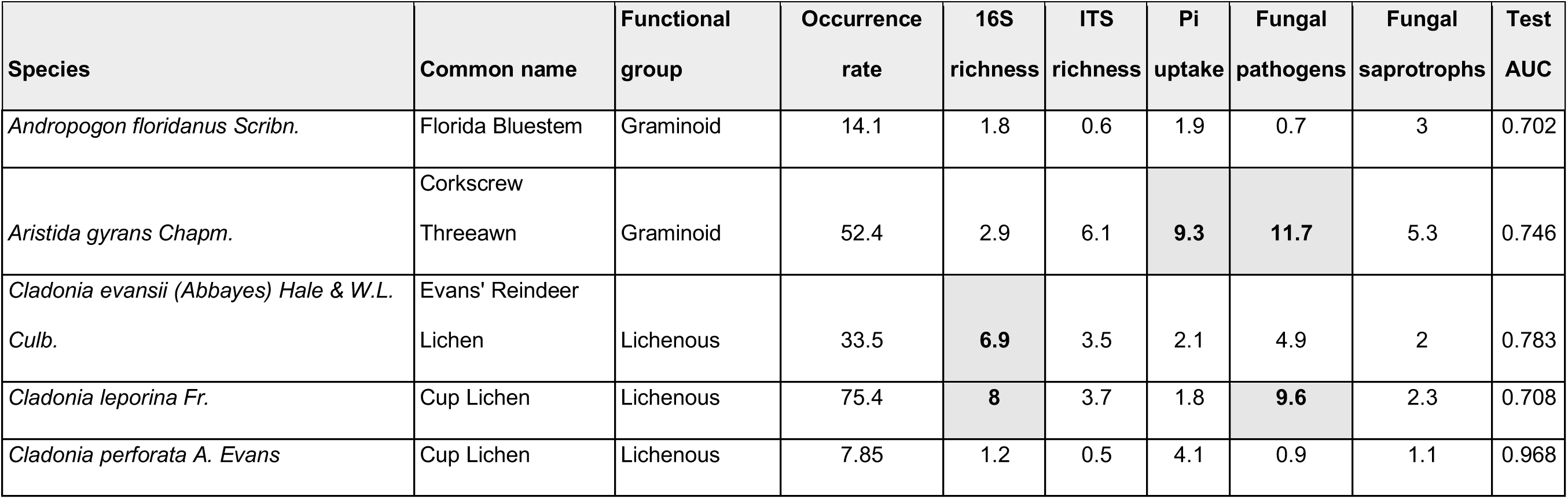
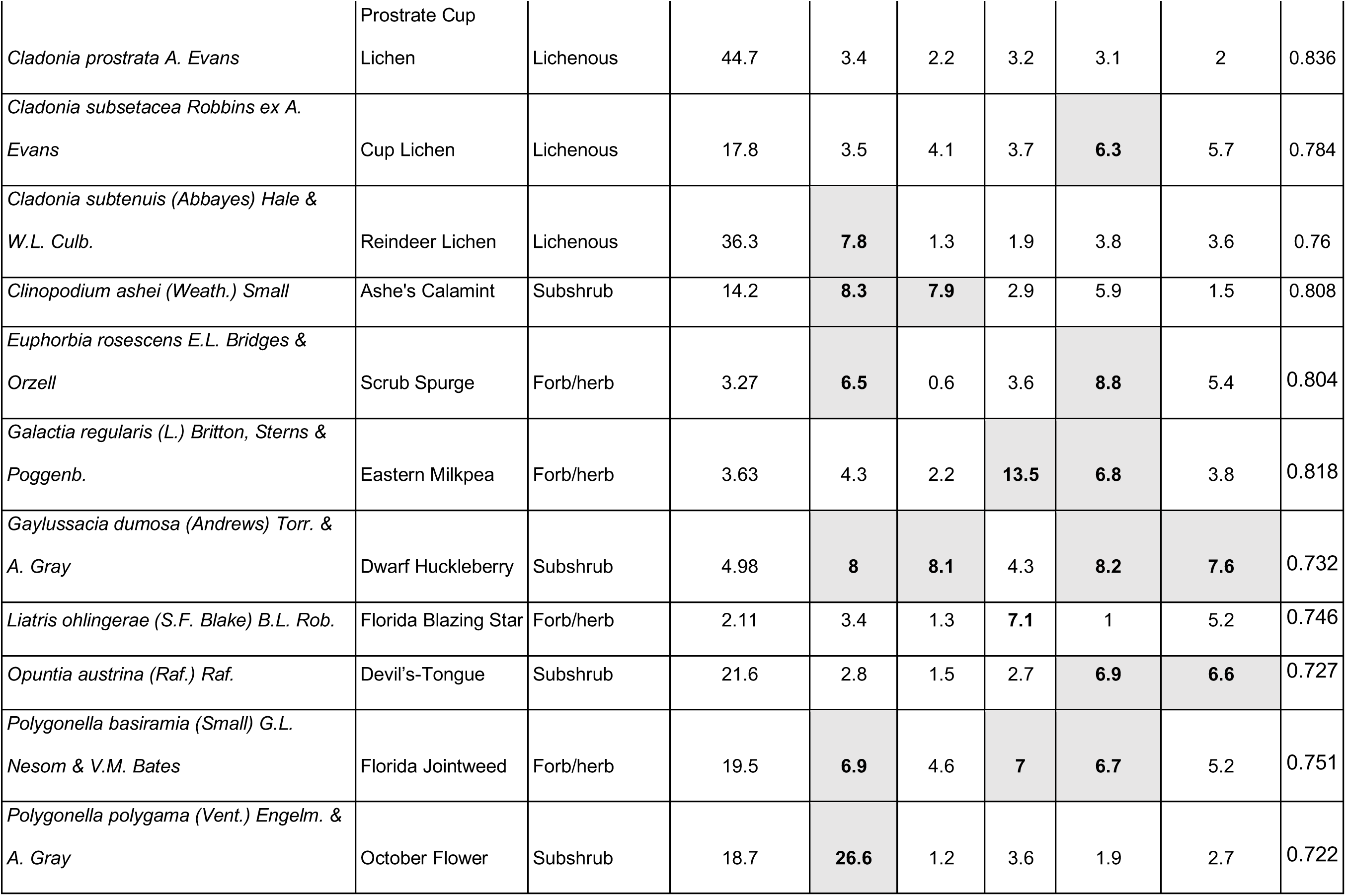
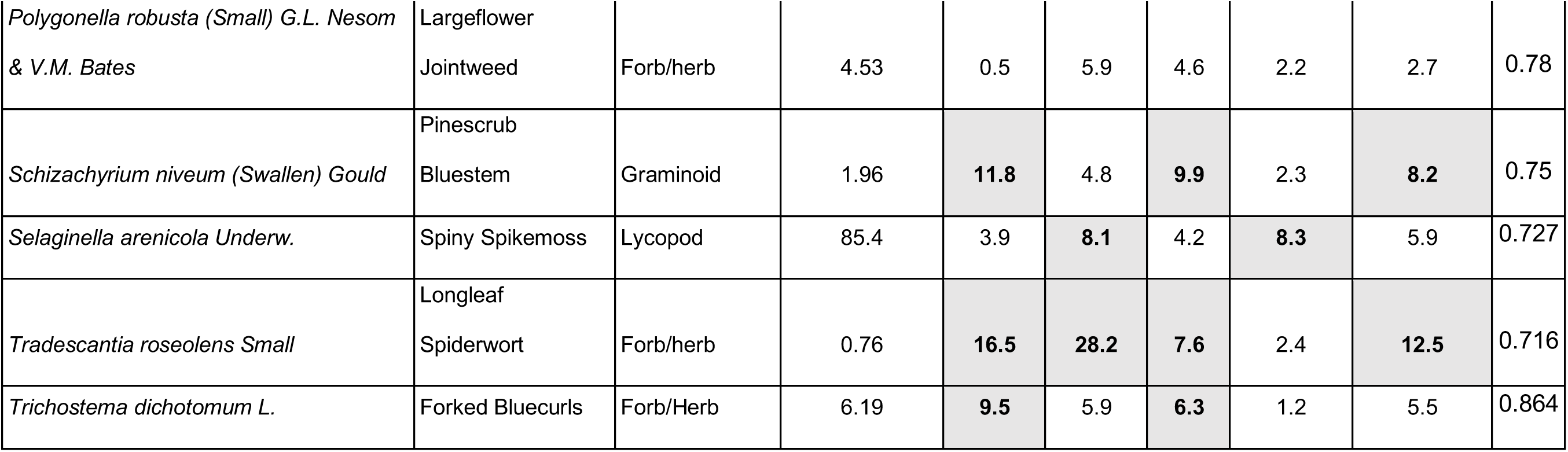
Variable importance (relative influence %) of the five microbial variables with the greatest explanatory power for 21 well-predicted (test AUC > 0.7) plant species distributions across the rosemary scrub landscape. Species and common name, as well as broad plant functional grouping (NRCS, 2025), and the occurrence rate across all sites (n = 676) are provided. Lichens were included in the plant community due to the large percentage of total ground cover they provide in this system. 16S richness indicates prokaryotic amplicon sequence variant richness, ITS richness indicates fungal operational taxonomic unit richness, Pi uptake indicates orthophosphate uptake/transfer gene abundance (predicted), fungal pathogens and saprotrophs indicate the relative abundance of all taxa classified in the respective trophic modes. Test AUC indicates total area under curve (AUC) for each plant species’ boosted regression tree model based on 10-fold cross validation. The values for each predictor variable with variable importance greater than expected by chance (>6.25%) are highlighted in bold.

### The soil microbiome and plant species distributions

Despite the generally strong effects of abiotic gradients on plant distributions, we identified six plant species whose distributions received >60% of their explanatory power from microbial predictor variables and 16 species for which microbial predictors accounted for >50% of occurrence probability (Figures 2-3). This suggests that for many plant species, the soil microbiome plays a critical role in structuring their presence on the landscape. Across the 21 informative SDMs, we found that prokaryotic richness, Pi transfer gene abundance, abundance of fungal pathogens, fungal richness, and abundance of fungal saprotrophs were the five most influential microbial predictors of species distributions (Table 1; Figures 2-3; Supplemental Figures 3-4). Prokaryotic richness had the greatest variable importance across plant species among microbial predictors (Table 1), and had greater importance than expected by chance for 11 of the 21 species distributions (Supplemental Figure 3), in one case (*Polygonella polygama*) accounting for 27% of species occurrence probability. Among these 11 species, eight occurred most often in sites with high prokaryotic richness, while the three species exhibiting the opposite trend were all lichens (Figure 4). Estimated abundance of prokaryotic phosphate transfer genes (*ugp*) was the second most important microbial predictor across models, accounting for more variance than expected by chance in 6 of the 21 plant species (Figure 2; Supplemental Figure 4). The distribution of the N-fixing legume *Galactia regularis* was strongly related to both abundance of *ugp* genes (13.5% variable importance) and nitrite reductase genes (*nir*; 10.6% variable importance), being found in open sand gaps with high values of *ugp* and extreme (both high and low) values of *nir*.

**Figure 3.**
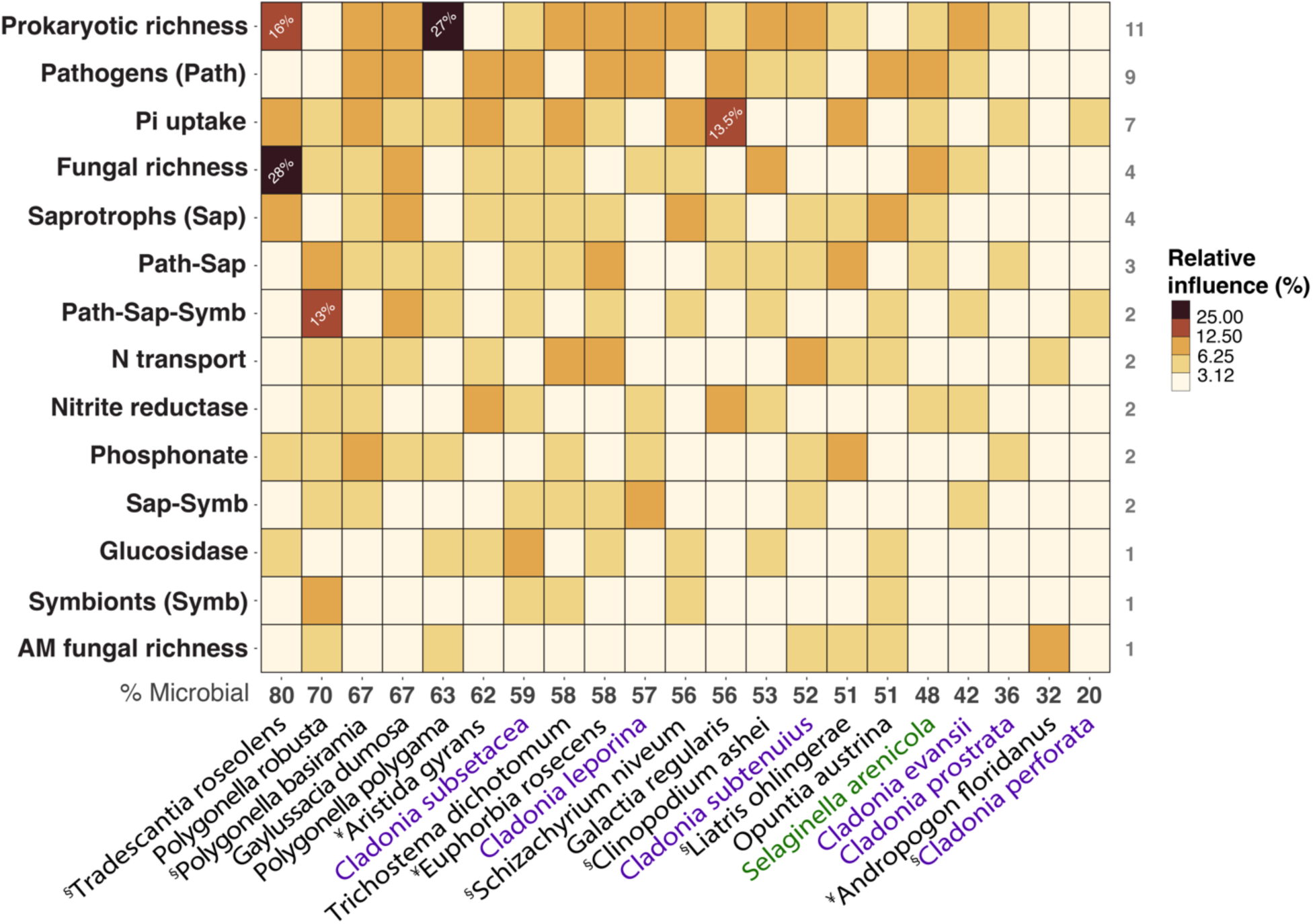
Heatmap of microbial predictor importance for all plant SDMs with test AUC > 0.70. The predictor variables are arranged along the y-axis in descending order of total relative influence across all plant SDMs. Counts of species for which each microbial predictor had a relative influence greater than expected by chance (>6.25%) are in gray (right), while values at the bottom indicate the total percentage of each species’ distribution explained by microbial metrics (% Microbial). For example, prokaryotic taxa richness had >6.25% influence for eleven plant SDMs, and microbial predictors provided 80% of the power in predicting *Tradescantia roseolens*’ distribution. The heatmap illustrates the microbial predictors that best explain plant distributions (top) and the plant species most responsive to these microbial factors (left). Lichens are in purple and the lycopod (*Selaginella arenicola*) is in green. ¥ = plant species endemic to Florida or southeastern USA. § = listed as a threatened or endangered plant species.

**Figure 4.**
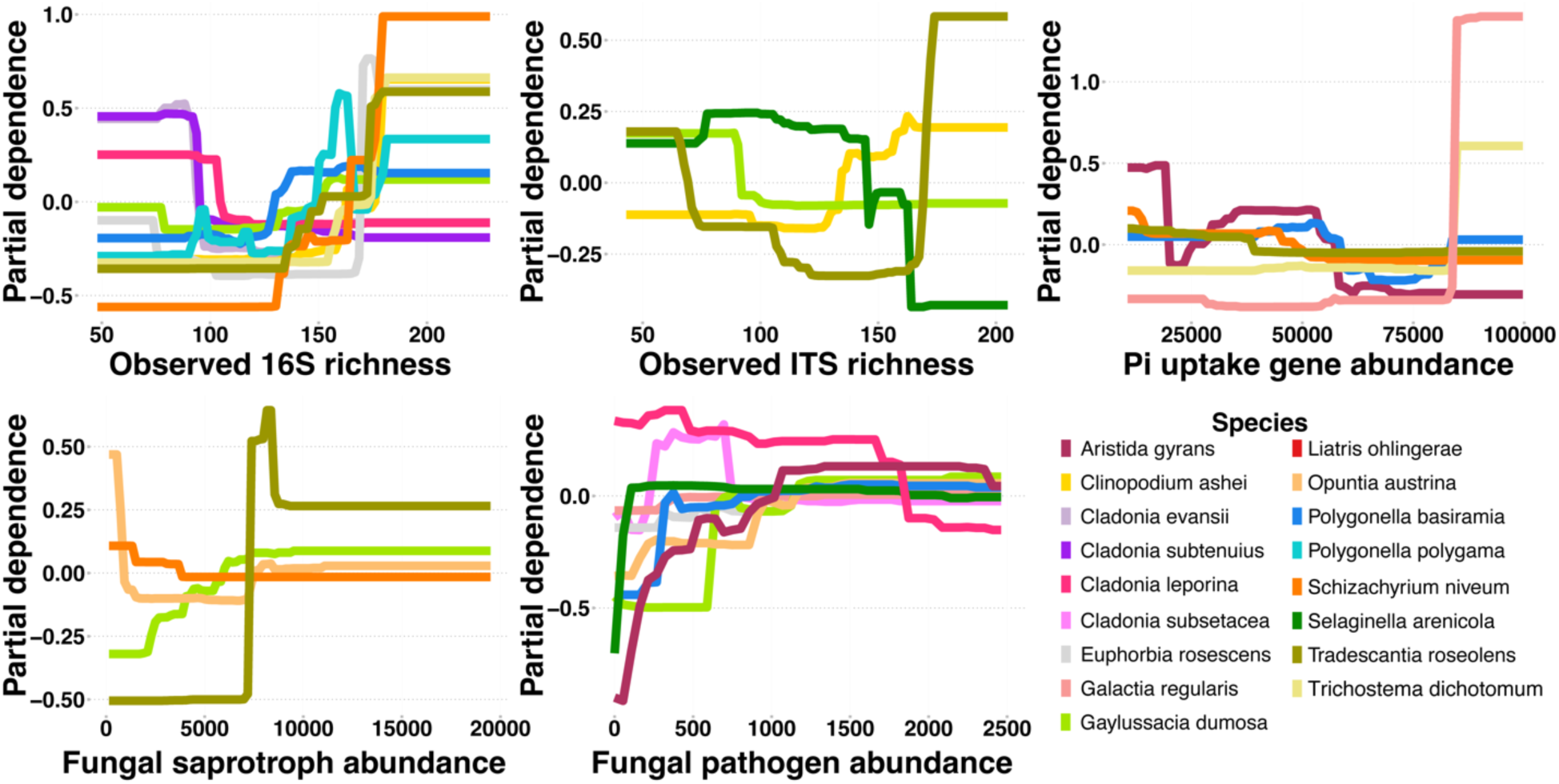
Partial dependence plots (scaled around zero to remove effects of overall abundance) for the five most explanatory microbial predictors across the set of plant species with well-explained distributions (test AUC > 0.70). Response curves are only shown for plant species whose variable importance was >6.25% (*i.e*., greater than expected by chance) for each predictor. Taxa whose predicted occurrence increases (higher partial dependence) along each x-axis indicates a positive relationship with each microbial predictor, while taxa whose predicted occurrence decreases along each x-axis indicates the reverse. Overall, plants tended to have a positive relationship with prokaryotic richness and, surprisingly, fungal pathogen abundance.

### Soil fungi and plant species distributions

The abundance of putative fungal pathogens was another particularly strong predictor across informative distribution models (influence greater than random expectation for nine of 21 species; Figure 3, Supplemental Figures 3-4). Somewhat surprisingly, six of these nine relationships were in the direction of plants being more common in open sand gaps with high pathogen loads. This positive association included all three of the endangered/endemic species that exhibited a strong response to fungal pathogens (*Aristida gyrans*, *Euphorbia rosescens*, and *Polygonella basiramia*). There was a noticeable difference in plants’ response to fungal pathogens based on functional group, with three non-vascular plants (two lichens and a lycopod) exhibiting distribution peaks at lower fungal pathogen abundance and six vascular plants (forbs, grasses and subshrubs) with distribution peaks associated with high fungal pathogen abundance. Of all fungal taxa classified as pathogenic (n = 38), we identified three taxa that could have played outsized roles in influencing species distributions through various mechanisms. *Curvularia lunata*, known to cause leaf spot in corn, was the most dominant taxon classified as a fungal pathogen in our study – with 94% frequency of occurrence across all sites and >40% relative fungal pathogen abundance per sample. *Gibberella fujikuroi*, a well-documented pathogen of rice and other monocots, was the second most dominant fungal pathogen. Thus, these dominant pathogens may play a particularly strong role in regulating grass species distributions, and the grass *Aristida gyrans* was the plant species that responded most strongly to fungal pathogen abundance (11.7% variable importance). The third most dominant fungal pathogen was *Lasiodiplodia crassispora*, a common pathogen on citrus plants, which may be present due to proximity of a major citrus agricultural zone. Finally, fungal richness was the other most important contributor to plant species distributions across the landscape. In particular, fungal richness was the most important predictor (28% variable influence) for the distribution of a threatened spiderwort, *Tradescantia roseolens* (Figure 3).

## Discussion

Following our predictions, we observed a particularly strong role of two abiotic gradients of known ecological importance in shaping the majority of plant species distributions across the landscape, but we also revealed that the soil microbiome played a previously unidentified and strong role in shaping the landscape distributions of many plants. As expected, TSF was the single greatest predictor for a majority of species distributions in our analysis (Figure 2). This variable was already known to be of great importance in shaping both plant and microbial communities in the Florida scrub habitat (Menges & Kohfeldt, 1995; Dee & Menges, 2014; Quintana-Ascencio *et al*., 2018) and is integral for the persistence of certain endangered plant species in this ecosystem (Menges & Hawkes, 1998; Quintana-ascencio *et al*., 2003; Quintana-Ascencio *et al*., 2018). Distance to the water table, a proxy for water availability, was another important environmental variable for species distributions in our study (Figure 2, Supplemental Figure 3). This metric, shaped by the topography of this ecosystem, has a clear role in filtering plants based on their rooting phenotypes (deep, taproot *versus* shallow, fibrous) and ability to survive according to natural water availability gradients (Abrahamson *et al*., 1984). It has also been shown that as elevation increases, not only does water availability decrease but so does nutrient availability (David *et al*., 2019), which suggests increased levels of stress for plants. Hernandez et al., (2018) showed that with these increased stresses, there is a significant reduction in microbial taxonomic richness, but also that the predominance of ‘positive interactions’ in soil microbial networks becomes greater. The strong influence of abiotic variables on plant distributions, as well as the effects of these variables on the soil microbiome, present potential problems in the future as climate change will alter fire regimes and water availability (Jones *et al*., 2022; Satoh *et al*., 2022), shifting locally adapted relationships between environment, plants, and the microbiome with unknown consequences for this unique and fragile ecosystem.

Plant species richness and many plant species distributions were similarly and strongly affected by four soil microbial metrics measured in our study, suggesting that specific components of the soil microbiome can shape plant communities across an entire landscape. These soil microbiome measures of landscape-scale importance included prokaryotic richness, fungal richness, abundance of fungal pathogens, and the abundance of prokaryotic phosphate transport genes. The abundance of fungal saprotrophs and nitrite reduction genes were also consistently important, but to a lesser degree than the previous four microbial properties (Figure 3, Supplemental Figure 3 & 4). Microbial richness in soil has been shown to contribute strongly to diversity and productivity in tallgrass prairies and to promote geochemical cycling (Vogelsang *et al*., 2006; Yuan *et al*., 2018), and is commonly linked to the maintenance of greater functional diversity, supporting a broader variety of ecosystem functions that can promote plant fitness (Bardgett & van der Putten, 2014; Delgado-Baquerizo *et al*., 2016). In our study, the distribution of a threatened plant species, *Tradescantia roseolens,* was the most responsive to microbial predictors overall (80%; Figure 2 & 3), and was strongly influenced by fungal richness (28%), prokaryotic richness (16%), and abundance of fungal saprotrophs (12%; Supplemental Figure 3). Eck *et al*., (2019) found that species-specific fungi have the capacity to influence plant germination and performance, thus shaping plant community dynamics, while Semchenko *et al*., (2018) suggest that plant-specific resource acquisition strategies (e.g., thin roots, high N concentrations) alter their interaction with different fungal guilds in soil and can result in negative feedbacks that impact community dynamics. Our results revealed that the threatened species, *T*. *roseolens* was highly responsive to soil fungal attributes (Figure 4), suggesting that maintenance of fungal diversity can be an important strategy for the preservation of threatened or endangered plants (Geisen *et al*., 2019).

The role of the soil microbiome in nutrient cycling is well-known, and multiple studies have identified important regulatory roles of microbial phosphorus cycling for plant health and productivity, typically in agricultural or restoration scenarios (Dai *et al*., 2018; Liang *et al*., 2020). Here we revealed that the abundance of prokaryotic phosphate transport genes (*ugp*) was a consistently influential microbial predictor of plant species distributions. These genes have been linked to increased plant-growth as well as plant disease resistance (Kang *et al*., 2024), which are both potential mechanisms that underpin the link between *ugp* genes and plant distributions, particularly for those taxa that have very strong relationships with lower or higher *ugp* abundance (Figure 4). To our knowledge, this is the first study to determine the predictive capacity of soil microbiome community attributes for plant species distributions, and presents an interesting path forward to determine whether a consistent picture emerges across diverse ecosystems regarding the importance of microbial richness, fungal pathogens, and microbe-driven nutrient cycling functions, or under what conditions different microbiome properties play key roles.

Our results also revealed shared responses to the abiotic gradients and microbial predictors that were shaped by membership in different plant groups, for example, rare versus common plants, non-vascular versus vascular plants, or leguminous versus non-leguminous plants. First, we observed that the rarer, endemic or threatened/endangered plant species generally responded more positively to higher abundance of fungal pathogens in soil than more abundant or common plant species. Many mechanisms of fungal pathogen control on plant diversity patterns have been summarized previously (see Bever *et al*., 2015), but our somewhat counter-intuitive result produces multiple potential explanations. First, rare plant species may benefit when fungal pathogens are abundant and attacking other more common, competitor plant species. This would suggest that fungal pathogen efficacy is species-specific and they are targeting more common plants. As part of this species-specificity, rare plant species may have developed some local pathogen resistance through modified root traits, immune system responses, or microbiome-mediation (Jones & Dangl, 2006; Vannier *et al*., 2019; Stump *et al*., 2020). An alternative explanation is that plant rarity itself could be due to fungal pathogens’ preferential attack on those species, a mechanism suggested by Kempel *et al*., (2018) where they found that the negative effects of soil fungal pathogens were two-times stronger for regionally rare plant species than regionally common plants. Along these lines, species-specific fungal pathogens have been shown to accumulate quickly on rare plant species, resulting in their lower densities, though whether pathogen accumulation is due to plant influence on the soil or fungal preference for their roots is not known (Klironomos, 2002). Another clear division between plant functional groups was the contrasting response of forbs and grasses (nine species) being most common at short TSF intervals due to the fact that many of these species maintain seed banks in soil that are triggered by fire or have higher germination and quicker growth rates (Menges & Hawkes, 1998; Navarra *et al*., 2011), while lichens (six species) were all more common after long TSF, likely as a result of lichens’ high mortality after surface fires, as well as their slowly-developing thalli, best-suited to later successional plant communities with some canopy cover (Menges & Kohfeldt, 1995; Yahr, 2000). Expanding phylogenetically on these functional groupings, we observed non-vascular plants (largely lichens) and vascular plants (largely forbs and grasses) having opposite responses to the soil microbiome, with non-vascular species responding negatively to prokaryotic richness while vascular plants preferred sites with greater prokaryotic richness (Figure 4). As non-vascular plants do not have true root systems (and lichens do not have any root-like structures), soil prokaryotic richness may simply increase in locations with more abundant vascular roots belowground, providing a diversity of carbon, preferential exudates, and nutrients sources (Steinauer *et al*., 2016; Curd *et al*., 2018; Liu *et al*., 2022). Alternatively, some studies have suggested that non-vascular plants can reduce microbial taxonomic richness by releasing certain secondary metabolites into their surrounding soil that hinder prokaryotic dispersal, growth, and survival in their proximity (Maier *et al*., 2014; Calcott *et al*., 2018; Prokopiev *et al*., 2025). Finally, we found that the distribution of leguminous *Galactia regularis* was positively related to the abundance of prokaryotic orthophosphate transfer and uptake (*ugp*) genes, which we suspect is the result of the high P demand to perform nitrogen-fixation (Nasto *et al*., 2014; Abdelrahman *et al*., 2018).

Beyond the interesting main effects of abiotic and microbial predictors on species distributions, there is also potential for these environmental factors to interact in their effect on plant distributions. Both TSF and fungal pathogens have strong effects on plant distributions, and their potential to interact is supported by previous experimental studies that found interactions between environmental disturbance and microbial-mediation of plant performance in this and other systems (Baynes *et al*., 2012; Revillini *et al*., 2022, 2023). For example, Revillini *et al*., (2022, 2023) demonstrated how common environmental stressors in the ecosystem directly altered the soil microbiome, which subsequently affected directionality of plant germination or productivity responses to those altered microbiomes. They also determined that the abundance of fungal pathogen *Gibberella fujikuroi* (O’Donnell *et al*., 2000) increased in response to both prescribed fire and allelochemical addition. In the current landscape-scale analysis, we identified *G. fujikuroi* as a dominant member of the fungal pathogen guild, a major microbial predictor of plant distributions (Figures 2-3). Given that the abundance of soil fungal pathogens can be enriched by environmental disturbances, and fungal pathogen abundance strongly altered plant distributions in the current study, important future research should first determine, definitively, whether these fungi are truly acting as pathogens on these plants, and if so, identify the mechanisms deployed by these pathogens to affect plant communities using explicit measures of microbial activity and function (*e.g.*, quantitative-stable isotope probing, metabolomics, transcriptomics). Following this approach, it would also be important to better define the role of shifting disturbance regimes in pathogen-mediation of plant health and species distributions as global changes progress. Further, the two predominant fungal pathogens in our study (*C. lunata* and *G. fujikuroi*) are commonly known to affect monocots, and we determined that the plant species responding most strongly to fungal pathogens was a regionally-endemic graminoid, *Aristida gyrans*, that was more common at sites with high fungal pathogen abundance (Figure 3). While we cannot determine causality in this relationship, and previous studies have provided evidence for a coinciding increase in fungal pathogen abundance with an increase in presence of preferentially-targeted plant species (Mommer *et al*., 2018), we posit the largely positive relationship observed between multiple plant species distributions and fungal pathogens in soil may result from competitive release, where the fungal suppression of targeted plant taxa opens niche space for those species whose distributions increased with fungal pathogens (Figure 4). More research explicitly investigating this potential effect should be conducted at similarly large scales in order to determine whether this relationship is a result of a build-up of fungal pathogen abundance due to the increasing presence of target plant species, or a plant-plant competitive release resulting from fungal pathogen impact on other plants.

Our results provide evidence for an alternative to the typical top-down perception of plant-microbial interactions and plant community development, where in response to their environment, plants selectively alter the composition and function of the soil microbiome (phytocentrism). While it is well-understood that plants and their associated microbiome interact through a series of environment-driven feedbacks between partners (Revillini *et al*., 2016; Teste *et al*., 2017; Beals *et al*., 2020), it is possible that the structure and function of the soil microbiome may play a previously unforeseen role in plant distributions regardless of abiotic conditions (Figure 1). Our analysis cannot establish causality, and there is always the possibility that a microbial property is associated with an unidentified environmental condition that is the true driver of plant distributions. However, after explicitly accounting for the abiotic gradients known to be most important in this system, there were multiple plant distributions (5 of 21 informative models) for which the influence of 2-4 microbial variables was equal to or greater than the influence of these two well-studied abiotic gradients (Figure 2 & Supplemental Figure 3), suggesting that the frequency of some plant species is critically dependent on properties of the soil microbiome. Our approach identified candidates for what the most important facets of the soil microbiome may be, and suggested a particularly strong role for microbial taxonomic richness and microbial nutrient-cycling in shaping landscape-level species distributions for approximately 38% of all plant species with informative models (Figure 2). Therefore, we explicitly call for international and legally-binding regulations that preserve or increase soil microbial diversity and focus on promoting beneficial soil functions such as water retention, carbon storage, and nutrient cycling, as in ‘<u>A Soil Deal for Europe’</u> (European Commission. Directorate General for Research and Innovation., 2023), but also for more tangible action to mitigate global changes that can generate negative feedbacks between below- and aboveground systems in the future.

## Supporting information

All Supporting Information

## Acknowledgments

D.R. received support from The Florida Native Plant Society (FNPS Research Grant). D.R., M.E.A. and C.A.S. were supported by NSF-DEB award 1922521. A.S.D. was supported by NSF-DEB award 2505582. B.K.A. was supported by the NSF-DGE Graduate Research Fellowship Program.

## Data Availability Statement

All metadata, and data processed for species distribution modeling as well as R script (for data analysis) are published in the figshare repository: DOI: 10.6084/m9.figshare.30156733. All 16S and ITS sequencing data are submitted to the NCBI SRA: XXIXIXIXI.

---

**Supplemental Figure 1.**
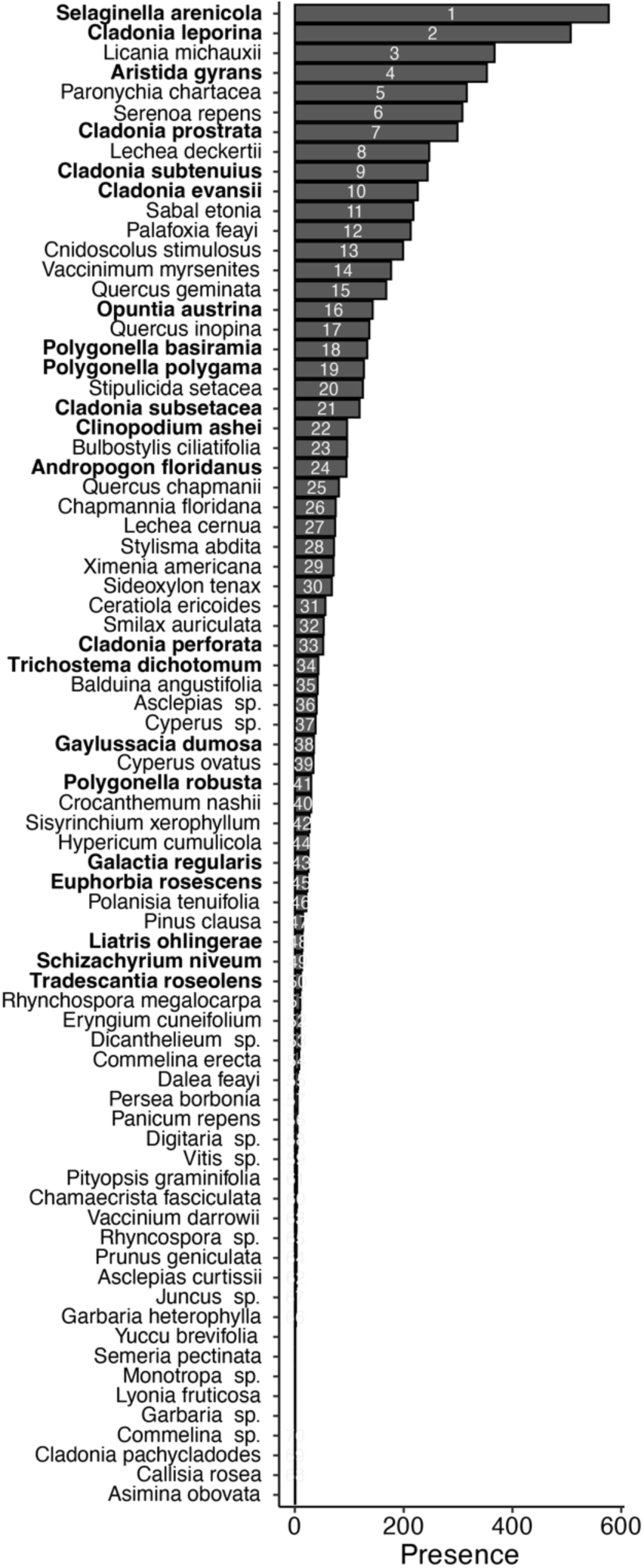
All plant species surveyed (n = 76) across all study sites (n = 676) organized according to their occurrence rate: presence across all study sites surveyed. Rank order is presented within bars. Species (n = 21) with informative species distribution models (test AUC > 0.7) are in bold.

**Supplemental Figure 2.**
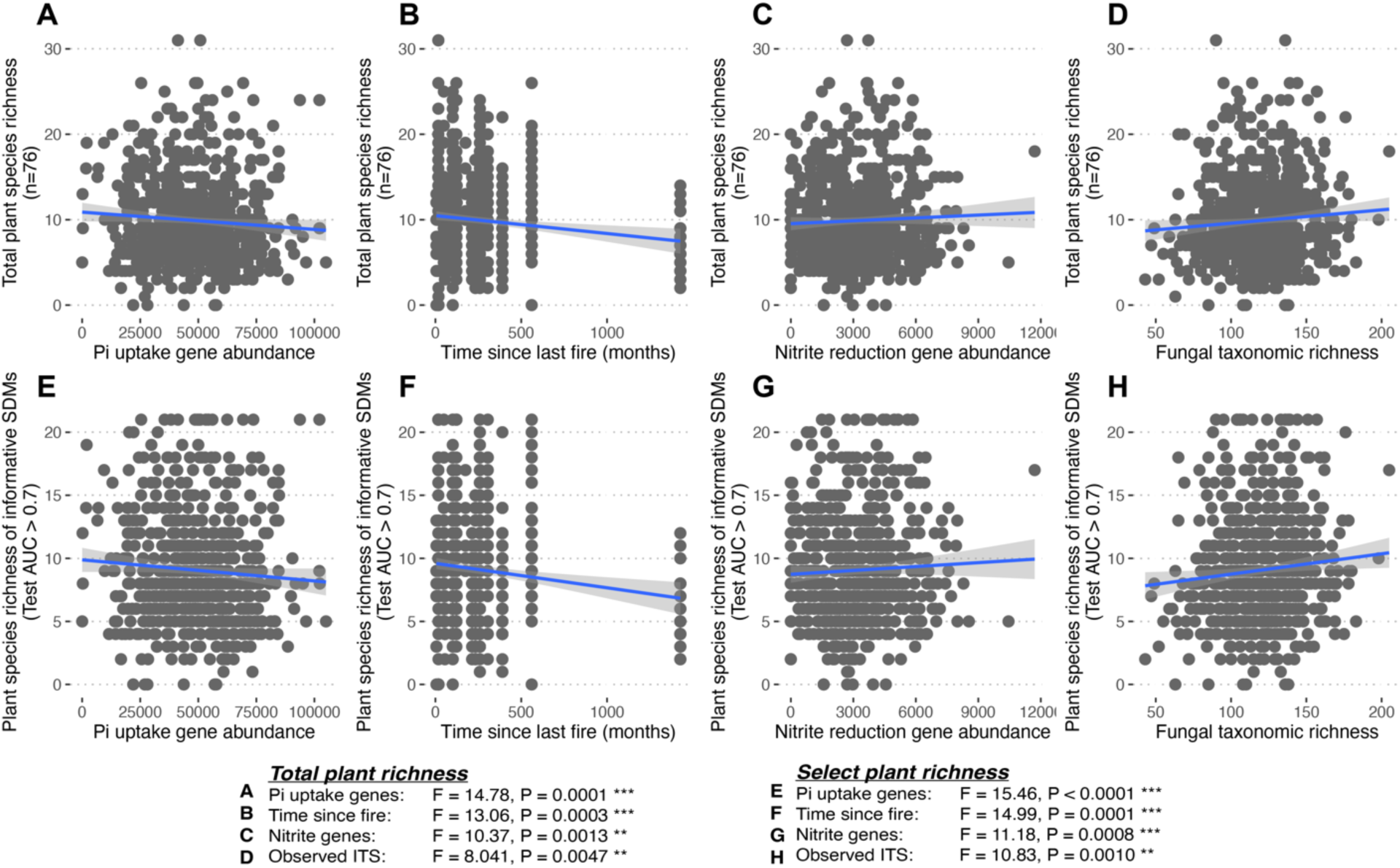
Significant relationships between abiotic and microbial predictors and plant richness across the landscape. Plant species richness of all identified species in our study (n = 76) as related to **A)** Pi uptake and transport gene (*ugp*) abundance, **B)** time since last fire, **C)** nitrite reducing gene abundance and **D)** observed taxonomic richness of fungi (ITS). Richness of plant species with informative SDMs (n = 21) as related to **E)** Pi uptake and transport gene (*ugp*) abundance, **F)** time since last fire, **G)** nitrite reducing gene abundance and **H)** observed taxonomic richness of fungi (ITS). Blue line represents linear correlation and shaded region represents 95% confidence interval. Statistical significance from generalized linear models generated by stepwise selection is presented for each figure on the bottom.

**Supplemental Figure 3.**
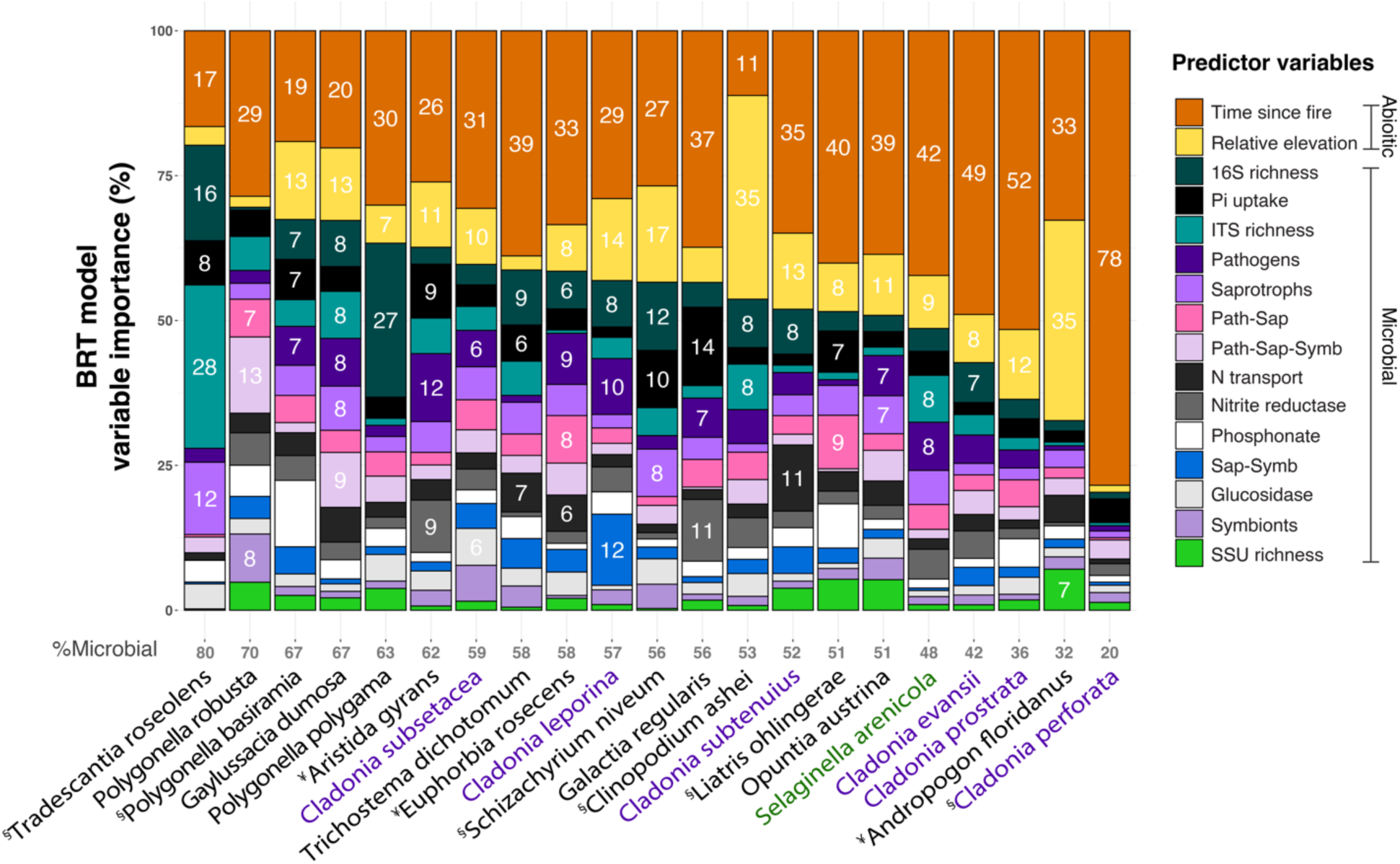
Total predictor importance for all plant SDMs with test AUC > 0.70, including abiotic variables time since last fire and relative elevation to the water table, as well as all microbial metrics. Total percent of microbial predictor influence on each species’ SDM is in gray (bottom). Plant species are arranged along the x-axis in descending order of total relative influence of microbial metrics. Lichens are in purple and the lycopod (*Selaginella arenicola*) is in green. ¥ = plant species endemic to Florida or southeastern USA. § = listed as a threatened or endangered plant species.

**Supplemental Figure 4.**
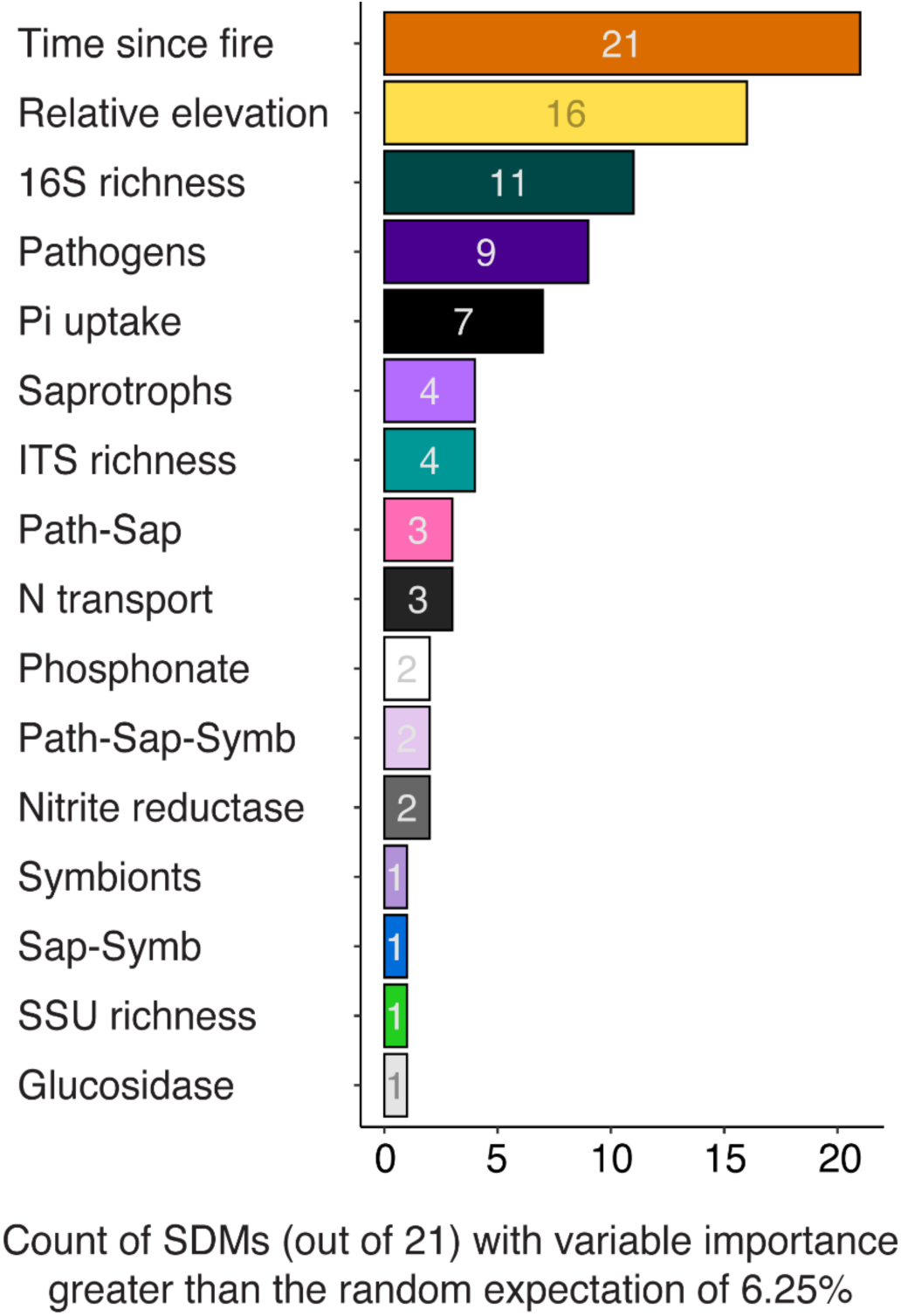
Summary of predictor variable impact on 21 plant species distribution models (SDMs) with test AUC > 0.70. Count of plant SDMs where the predictor explained greater than the random expectation of 6.25%. For example, time since last fire had a greater impact on 100% of the SDMs than expected by chance, and prokaryotic (16S) species richness had a greater impact on 52% of the SDMs than expected by chance.

**Supplemental Figure 5.**
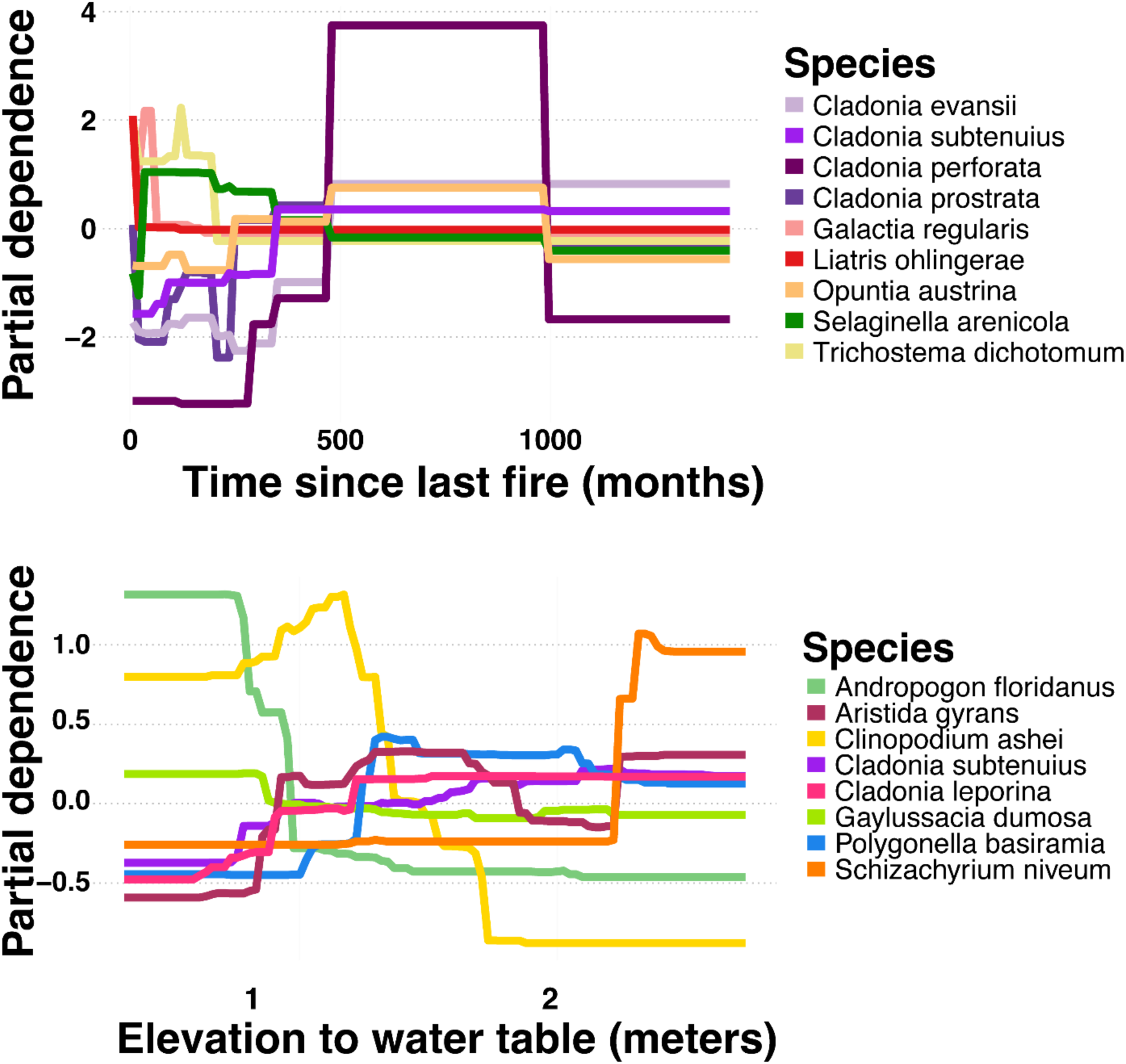
Partial dependence plots (scaled to a mean of zero to remove effects of overall abundance) for the two abiotic predictors: time since last fire (months) and relative elevation to the water table (meters). Only plant species whose time since last fire variable importance was >35% (mean) and relative elevation variable importance was >11.2% (mean) are shown to emphasize plant taxa with distributions strongly predicted by these abiotic variables. Plant species exhibited a fairly even mix of responses to these abiotic gradients, demonstrating specialization on different parts of these niche axes.

**Supplemental Figure 6.**
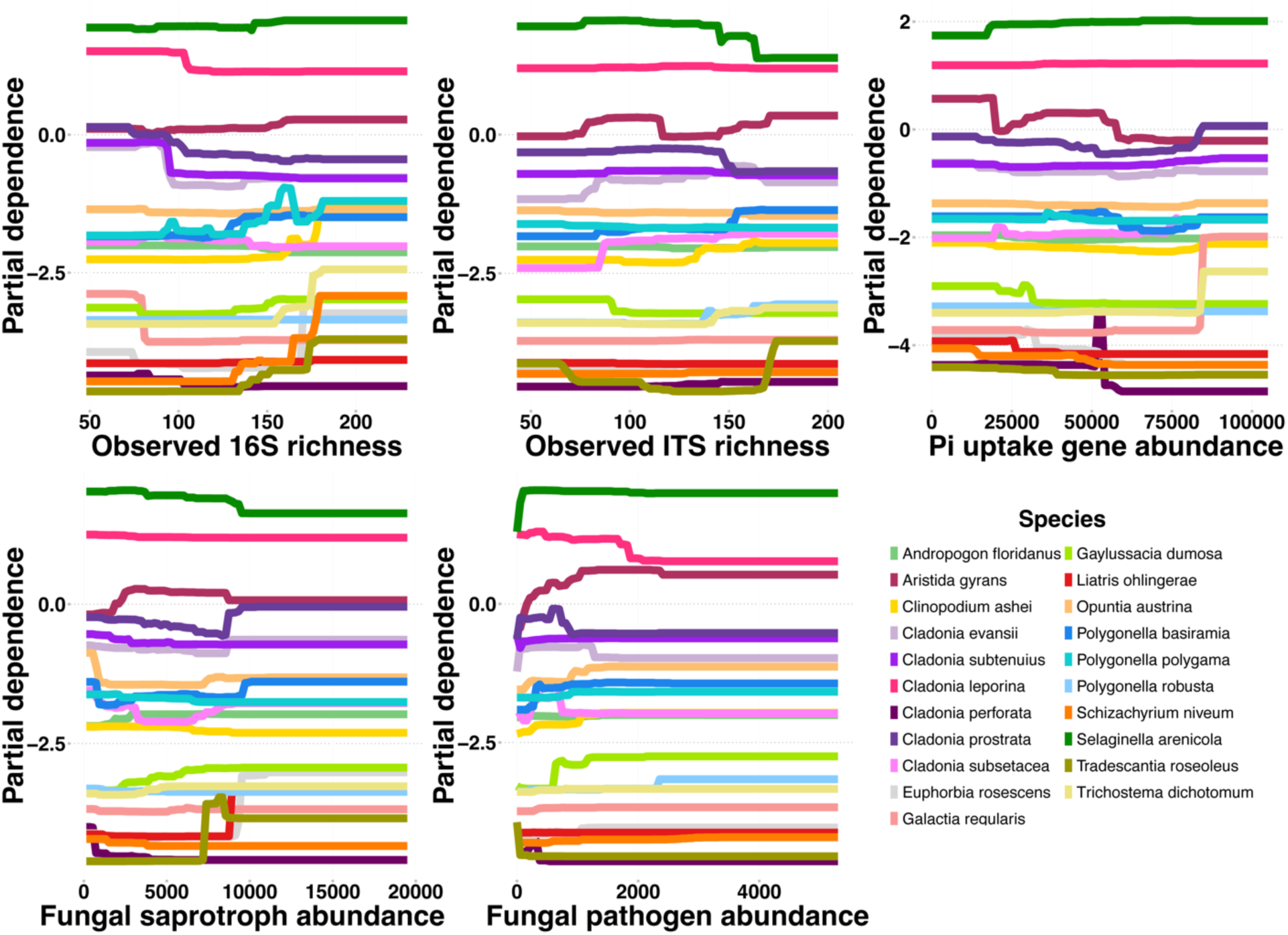
Partial dependence plots for the five most explanatory microbial predictor variables across the 21 species distribution models with test AUC > 0.70.

## References

Abdelrahman M, El-Sayed MA, Hashem A, Abd-Allah EF, Alqarawi AA, Burritt DJ, Tran LSP. 2018. Metabolomics and transcriptomics in legumes under phosphate deficiency in relation to nitrogen fixation by root nodules. Frontiers in Plant Science 9: 1–8.

Abrahamson WG, Johnson AF, Layne JN, Peroni PA. 1984. Vegetation of Archbold Station, Florida: An example of the southern Lake Wales Ridge. Florida Scientist 47: 209–250.

Afkhami ME. 2023. Past microbial stress benefits tree resilience. Science 380: 798–799.

Afkhami ME, Almeida BK, Hernandez DJ, Kiesewetter KN, Revillini DP. 2020. Tripartite mutualisms as models for understanding plant–microbial interactions. Current Opinion in Plant Biology 56: 28–36.

Afkhami ME, Mcintyre PJ, Strauss SY. 2014. Mutualist-mediated effects on species’ range limits across large geographic scales. Ecology Letters 17: 1265–1273.

Allsup CM, George I, Lankau RA. 2023. Shifting microbial communities can enhance tree tolerance to changing climates. Science 380: 835–840.

Anthony MA, Bender SF, van der Heijden MGA. 2023. Enumerating soil biodiversity. Proceedings of the National Academy of Sciences 120: e2304663120.

Bagchi R, Gallery RE, Gripenberg S, Gurr SJ, Narayan L, Addis CE, Freckleton RP, Lewis OT. 2014. Pathogens and insect herbivores drive rainforest plant diversity and composition. Nature 506: 85–8.

Bardgett RD, van der Putten WH. 2014. Belowground biodiversity and ecosystem functioning. Nature 515: 505–511.

Baynes M, Newcombe G, Dixon L, Castlebury L, O’Donnell K. 2012. A novel plant-fungal mutualism associated with fire. Fungal Biology 116: 133–144.

Beals KK, Moore JAM, Kivlin SN, Bayliss SLJ, Lumibao CY, Moorhead LC, Patel M, Summers JL, Ware IM, Bailey JK, et al. 2020. Predicting Plant-Soil Feedback in the Field: Meta-Analysis Reveals That Competition and Environmental Stress Differentially Influence PSF. Frontiers in Ecology and Evolution 8: 191.

Bever JD. 2002. Negative feedback within a mutualism: host-specific growth of mycorrhizal fungi reduces plant benefit. Proceedings of the Royal Society of Britain Biological sciences 269: 2595–2601.

Bever JD, Mangan SA, Alexander HM. 2015. Maintenance of Plant Species Diversity by Pathogens. *Annual Review of Ecology*, Evolution, and Systematics 46: 305–325.

Blonder B. 2018. Hypervolume concepts in niche- and trait-based ecology. Ecography 41: 1441–1455.

Bolyen E, Rideout JR, Dillon MR, Bokulich NA, Abnet CC, Al-ghalith GA, Alexander H, Alm EJ, Arumugam M, Asnicar F, et al. 2019. Reproducible, interactive, scalable and extensible microbiome data science using QIIME 2. Nature Biotechnology 37: 852–857.

Calcott MJ, Ackerley DF, Knight A, Keyzers RA, Owen JG. 2018. Secondary metabolism in the lichen symbiosis. Chemical Society Reviews 47: 1730–1760.

Callahan BJ, McMurdie PJ, Rosen MJ, Han AW, Johnson AJA, Holmes SP. 2016. DADA2: High-resolution sample inference from Illumina amplicon data. Nature Methods 13: 581–583.

Callaway RM, Thelen GC, Rodriguez A, Holben WE. 2004. Soil biota and exotic plant invasion. Nature 427: 731–733.

Curd EE, Martiny JBH, Li H, Smith TB. 2018. Bacterial diversity is positively correlated with soil heterogeneity. Ecosphere 9: e02079.

Dai Z, Su W, Chen H, Barberán A, Zhao H, Yu M, Yu L, Brookes PC, Schadt CW, Chang SX, et al. 2018. Long-term nitrogen fertilization decreases bacterial diversity and favors the growth of Actinobacteria and Proteobacteria in agro-ecosystems across the globe. Global Change Biology 24: 3452–3461.

David AS, Quintana-Ascencio PF, Menges ES, Thapa-Magar KB, Afkhami ME, Searcy CA. 2019. Soil microbiomes underlie population persistence of an endangered plant species. American Naturalist 194: 488–494.

David AS, Thapa-Magar KB, Afkhami ME. 2018. Microbial mitigation–exacerbation continuum: a novel framework for microbiome effects on hosts in the face of stress. Ecology 99: 517–523.

De Vries FT, Griffiths RI, Knight CG, Nicolitch O, Williams A. 2020. Harnessing rhizosphere microbiomes for drought-resilient crop production. Science 368: 270–274.

De Vries FT, Liiri ME, Bjørnlund L, Bowker MA, Christensen SS, Setälä HM, Bardgett RD, Bjornlund L, Bowker MA, Christensen SS, et al. 2012. Land use alters the resistance and resilience of soil food webs to drought. Nature Climate Change 2: 276–280.

Dean S. 2015. Fire Effects on Soil Biogeochemistry in Florida Scrubby Flatwoods. The American Midland Naturalist 174: 49–64.

Dee JR, Menges ES. 2014. Gap ecology in the Florida scrubby flatwoods: Effects of time-since-fire, gap area, gap aggregation and microhabitat on gap species diversity. Journal of Vegetation Science 25: 1235–1246.

Delavaux CS, Crowther TW, Bever JD, Weigelt P, Gora EM. 2024. Mutualisms weaken the latitudinal diversity gradient among oceanic islands. Nature 627: 335–339.

Delgado-Baquerizo M, Maestre FT, Reich PB, Jeffries TC, Gaitan JJ, Encinar D, Berdugo M, Campbell CD, Singh BK. 2016. Microbial diversity drives multifunctionality in terrestrial ecosystems. Nature Communications 7: 1–8.

Delgado-Baquerizo M, Reich PB, Trivedi C, Eldridge DJ, Abades S, Alfaro FD, Bastida F, Berhe AA, Cutler NA, Gallardo A, et al. 2020. Multiple elements of soil biodiversity drive ecosystem functions across biomes. Nature Ecology and Evolution 4: 210–220.

Dobson A, Crawley M. 1994. Pathogens and the structure of plant communities. Trends in Ecology & Evolution 9: 393–398.

Douglas GM, Maffei VJ, Zaneveld JR, Yurgel SN, Brown JR, Taylor CM, Huttenhower C, Langille MGI. 2020. PICRUSt2 for prediction of metagenome functions. Nature Biotechnology 38: 669–673.

Eck JL, Stump SM, Delavaux CS, Mangan SA, Comita LS. 2019. Evidence of within-species specialization by soil microbes and the implications for plant community diversity. Proceedings of the National Academy of Sciences of the United States of America 116: 7371–7376.

Elith J, Leathwick JR, Hastie T. 2008. A working guide to boosted regression trees. Journal of Animal Ecology 77: 802–813.

European Commission. Directorate General for Research and Innovation. 2023. EU missions: soil deal for Europe : what is the EU mission : a soil deal for Europe. LU: Publications Office.

Fischer NH, Williamson GB, Weidenhamer JD, Richardson DR. 1994. In search of allelopathy in the Florida scrub: The role of terpenoids. Journal of Chemical Ecology 20: 1355– 1380.

Friedman JH. 2002. Stochastic gradient boosting. Computational Statistics & Data Analysis 38: 367–378.

García-Palacios P, Maestre FT, Bardgett RD, de Kroon H. 2012. Plant responses to soil heterogeneity and global environmental change. Journal of Ecology 100: 1303–1314.

Geisen S, Wall DH, van der Putten WH. 2019. Challenges and Opportunities for Soil Biodiversity in the Anthropocene. Current Biology 29: R1036–R1044.

Gohl DM, Vangay P, Garbe J, MacLean A, Hauge A, Becker A, Gould TJ, Clayton JB, Johnson TJ, Hunter R, et al. 2016. Systematic improvement of amplicon marker gene methods for increased accuracy in microbiome studies. Nature Biotechnology 34: 942–949.

van der Heijden MGA, Bardgett RD, van Straalen NM. 2008. The unseen majority: soil microbes as drivers of plant diversity and productivity in terrestrial ecosystems. Ecology Letters 11: 296–310.

Hernandez DJ, David AS, Menges ES, Searcy CA, Afkhami ME. 2021. Environmental stress destabilizes microbial networks. ISME Journal 15: 1722–1734.

Hijmans RJ, Phillips S, Leathwick J, Elith J. 2024. dismo: Species Distribution Modeling.

Jansson JK, Hofmockel KS. 2020. Soil microbiomes and climate change. Nature Reviews Microbiology 18: 35–46.

Jones MW, Abatzoglou JT, Veraverbeke S, Andela N, Lasslop G, Forkel M, Smith AJP, Burton C, Betts RA, van der Werf GR, et al. 2022. Global and Regional Trends and Drivers of Fire Under Climate Change. Reviews of Geophysics 60: 1–76.

Jones JDG, Dangl JL. 2006. The plant immune system. Nature 444: 323–329.

Kandlikar GS, Johnson CA, Yan X, Kraft NJB, Levine JM. 2019. Winning and losing with microbes: how microbially mediated fitness differences influence plant diversity. Ecology Letters 22: 1178–1191.

Kang L, Song Y, Mackelprang R, Zhang D, Qin S, Chen L, Wu L, Peng Y, Yang Y. 2024. Metagenomic insights into microbial community structure and metabolism in alpine permafrost on the Tibetan Plateau. Nature Communications 15: 5920.

Katoh K, Standley DM. 2013. MAFFT multiple sequence alignment software version 7: Improvements in performance and usability. Molecular Biology and Evolution 30: 772–780.

Kempel A, Rindisbacher A, Fischer M, Allan E. 2018. Plant soil feedback strength in relation to large-scale plant rarity and phylogenetic relatedness. Ecology 99: 597–606.

Kiers ET, Denison RF. 2008. Sanctions, Cooperation and the Stability of Plant-Rhizosphere Mutualisms. Annual Review of Ecology, Evolution, and Systematics 39: 215–236.

Klironomos JN. 2002. Feedback with soil biota contributes to plant rarity and invasiveness in communities. Nature 417: 67–70.

Liang JL, Liu J, Jia P, Yang T tao, Zeng Q wei, Zhang S chang, Liao B, Shu W sheng, Li J tian. 2020. Novel phosphate-solubilizing bacteria enhance soil phosphorus cycling following ecological restoration of land degraded by mining. ISME Journal 14: 1600–1613.

Liu H, Brettell LE, Qiu Z, Singh BK. 2020. Microbiome-Mediated Stress Resistance in Plants. Trends in Plant Science 25: 733–743.

Liu B, Han F, Ning P, Li H, Rengel Z. 2022. Root traits and soil nutrient and carbon availability drive soil microbial diversity and composition in a northern temperate forest. Plant and Soil 479: 281–299.

Lugtenberg B, Kamilova F. 2009. Plant-growth-promoting rhizobacteria. Annual Review of Microbiology 63: 541–56.

Maier S, Schmidt TSB, Zheng L, Peer T, Wagner V, Grube M. 2014. Analyses of dryland biological soil crusts highlight lichens as an important regulator of microbial communities. Biodiversity and Conservation 23: 1735–1755.

Mangan SA, Schnitzer SA, Herre EA, Mack KML, Valencia MC, Sanchez EI, Bever JD. 2010. Negative plant–soil feedback predicts tree-species relative abundance in a tropical forest. Nature 466: 752–755.

Menges ES, Craddock A, Salo J, Zinthefer R, Weekley CW. 2008. Gap ecology in Florida scrub: Species occurrence, diversity and gap properties. Journal of Vegetation Science 19: 503–514.

Menges ES, Hawkes CV. 1998. Interactive Effects of Fire and Microhabitat on Plants of Florida Scrub. Ecological Applications 8: 935–946.

Menges ES, Kohfeldt N. 1995. Life History Strategies of Florida Scrub Plants in Relation to Fire. Bulletin of the Torrey Botanical Club 122: 282–297.

Menges ES, Main KN, Pickert RL, Ewing K. 2017. Evaluating a fire management plan for fire regime goals in a Florida landscape. Natural Areas Journal 37: 212–227.

Mills KE, Bever JD. 1998. Maintenance of diversity within plant communities: Soil pathogens as agents of negative feedback. Ecology 79: 1595–1601.

Mommer L, Cotton TEA, Raaijmakers JM, Termorshuizen AJ, van Ruijven J, Hendriks M, van Rijssel SQ, van de Mortel JE, van der Paauw JW, Schijlen EGWM, et al. 2018. Lost in diversity: the interactions between soil-borne fungi, biodiversity and plant productivity. New Phytologist 218: 542–553.

Nasto MK, Alvarez-Clare S, Lekberg Y, Sullivan BW, Townsend AR, Cleveland CC. 2014. Interactions among nitrogen fixation and soil phosphorus acquisition strategies in lowland tropical rain forests. Ecology Letters 17: 1282–1289.

Navarra JJ, Kohfeldt N, Menges ES, Quintana-Ascencio PF. 2011. Seed bank changes with time-since-fire in Florida rosemary scrub. Fire Ecology 7: 17–31.

Neu AT, Allen EE, Roy K. 2021. Defining and quantifying the core microbiome: Challenges and prospects. PNAS 118: e2104429118.

Nguyen NH, Song Z, Bates ST, Branco S, Tedersoo L, Menke J, Schilling JS, Kennedy PG. 2016. FUNGuild: An open annotation tool for parsing fungal community datasets by ecological guild. Fungal Ecology 20: 241–248.

O’Donnell K, Nirenberg HI, Aoki T, Cigelnik E. 2000. A multigene phylogeny of the Gibberella fujikuroi species complex. Mycoscience 41: 61–78.

Peay KG. 2016. The Mutualistic Niche: Mycorrhizal Symbiosis and Community Dynamics. *Annual Review of Ecology*, Evolution, and Systematics 47: 143–164.

Prokopiev IA, Sazanova KV, Sleptsov IV, Filippova GV, Kuzmina NP, Frolova DA, Zholobova ZhO. 2025. Effect of Secondary Metabolites of Lichens on Microbial Communities in Permafrost Forest Soils. Contemporary Problems of Ecology 18: 82–100.

Qu X, Li X, Bardgett RD, Kuzyakov Y, Revillini D, Sonne C, Xia C, Ruan H, Liu Y, Cao F, et al. 2024. Deforestation impacts soil biodiversity and ecosystem services worldwide. Proceedings of the National Academy of Sciences 121: e2318475121.

Quintana-Ascencio PF, Koontz SM, Smith SA, Sclater VL, David AS, Menges ES. 2018. Predicting landscape-level distribution and abundance: Integrating demography, fire, elevation and landscape habitat configuration. Journal of Ecology 106: 2395–2408.

Quintana-ascencio PF, Menges ES, Weekley CW. 2003. A Fire-Explicit Population Viability Analysis of Hypericum cumulicola in Florida Rosemary Scrub. Conservation Biology 17: 433– 449.

R Core Team. 2020. R: A language and environment for statistical computing.

Revillini D, David AS, Menges ES, Main KN, Afkhami ME, Searcy CA. 2022. Microbiome-mediated response to pulse fire disturbance outweighs the effects of fire legacy on plant performance. New Phytologist 233: 2071–2082.

Revillini D, David AS, Reyes AL, Knecht LD, Vigo C, Allen P, Searcy CA, Afkhami ME. 2023. Allelopathy-selected microbiomes mitigate chemical inhibition of plant performance. New Phytologist 240: 2007–2019.

Revillini D, Gehring CA, Johnson NC. 2016. The role of locally adapted mycorrhizas and rhizobacteria in plant-soil feedback systems. Functional Ecology 30: 1086–1098.

Rillig MC. 2004. Arbuscular mycorrhizae and terrestrial ecosystem processes. Ecology Letters 7: 740–754.

Rudgers JA, Gehring CA, Taylor DL, Taylor MD, Chung YA. 2025. Integration of plant–soil feedbacks with resilience theory for climate change. Trends in Ecology & Evolution 40: 749– 759.

Satoh Y, Yoshimura K, Pokhrel Y, Kim H, Shiogama H, Yokohata T, Hanasaki N, Wada Y, Burek P, Byers E, et al. 2022. The timing of unprecedented hydrological drought under climate change. Nature Communications 13: 3287.

Sayer EJ, Oliver AE, Fridley JD, Askew AP, Mills RTE, Grime JP. 2017. Links between soil microbial communities and plant traits in a species-rich grassland under long-term climate change. Ecology and Evolution 7: 855–862.

Schnitzer SA, Klironomos JN, HilleRisLambers J, Kinkel LL, Reich PB, Xiao K, Rillig MC, Sikes BA, Callaway RM, Mangan SA, et al. 2011. Soil microbes drive the classic plant diversity-productivity pattern. Ecology 92: 296–303.

Semchenko M, Leff JW, Lozano YM, Saar S, Davison J, Wilkinson A, Jackson BG, Pritchard WJ, De Long JR, Oakley S, et al. 2018. Fungal diversity regulates plant-soil feedbacks in temperate grassland. Science Advances 4: eaau4578.

Steinauer K, Chatzinotas A, Eisenhauer N. 2016. Root exudate cocktails: the link between plant diversity and soil microorganisms? Ecology and Evolution 6: 7387–7396.

Stump SM, Marden JH, Beckman NG, Mangan SA, Comita LS. 2020. Resistance genes affect how pathogens maintain plant abundance and diversity. American Naturalist 196: 472– 486.

Taylor BN, Simms EL, Komatsu KJ. 2020. More than a functional group: Diversity within the legume-rhizobia mutualism and its relationship with ecosystem function. Diversity 12: 50.

Teste FP, Kardol P, Turner BL, Wardle DA, Zemunik G, Renton M, Laliberté E. 2017. Plant-soil feedback and the maintenance of diversity in Mediterranean-climate shrublands. Science 355: 173–176.

Trivedi P, Delgado-Baquerizo M, Trivedi C, Hu H, Anderson IC, Jeffries TC, Zhou J, Singh BK. 2016. Microbial regulation of the soil carbon cycle: Evidence from gene-enzyme relationships. ISME Journal 10: 2593–2604.

Trivedi P, Leach JE, Tringe SG, Sa T, Singh BK. 2020. Plant–microbiome interactions: from community assembly to plant health. Nature Reviews Microbiology 18: 607–621.

Valavi R, Guillera-Arroita G, Lahoz-Monfort JJ, Elith J. 2022. Predictive performance of presence-only species distribution models: a benchmark study with reproducible code. Ecological Monographs 92: e01486.

Vannier N, Agler M, Hacquard S. 2019. Microbiota-mediated disease resistance in plants. PLoS Pathogens 15: 1–7.

Vogelsang KM, Reynolds HL, Bever JD. 2006. Mycorrhizal fungal identity and richness determine the diversity and productivity of a tallgrass prairie system. New Phytologist 172: 554– 562.

Wagg C, Schlaeppi K, Banerjee S, Kuramae EE, van der Heijden MGA. 2019. Fungal-bacterial diversity and microbiome complexity predict ecosystem functioning. Nature communications 10: 4841.

Westover KM, Bever JD, Carolina N. 2001. Mechanisms of Plant Species Coexistence: Roles of Rhizosphere Bacteria and Root Fungal Pathogens. Ecology 82: 3285–3294.

Yahr R. 2000. Ecology and post-fire recovery of Cladonia perforata, an endangered Florida-scrub lichen. Forest Snow and Landscape Research. [print] 2000; 75: 339–356.

Yang G, Wagg C, Veresoglou SD, Hempel S, Rillig MC. 2018. How Soil Biota Drive Ecosystem Stability. Trends in Plant Science 23: 1057–1067.

Yuan X, Knelman JE, Wang D, Goebl A, Gasarch E, Yuan X, Knelman JE, Wang D, Goebl A, Gasarch E, et al. 2018. Patterns of Soil Bacterial Richness and Composition Tied to Plant Richness, Soil Nitrogen, and Soil Acidity in Alpine Tundra. Arctic, Antarctic, and Alpine Research 49: 441–453.

Zhang T, Chen HYH, Ruan H. 2018. Global negative effects of nitrogen deposition on soil microbes. ISME Journal 12: 1817–1825.

