## Supplementary material for "The microbial landscape: soil microbiome properties predict plant species distributions": All Supporting Information

### Detailed Methods

Adapted protocol for DNeasy PowerSoil Pro QIAcube HT Kit without QiaCube robot (Qiagen, Carlsbad, CA, USA). Briefly, we used ~550 mg of soil for extractions, performed all wash and elution steps using S-Blocks (Cat. ID 19585) in an Avanti JXN-26 centrifuge (Beckman Coulter Inc., Brea, CA, USA) at 4500 RCF (G-force), and performed final elution with 50 µL 10 mM Tris-HCl.

### Supplemental Figures

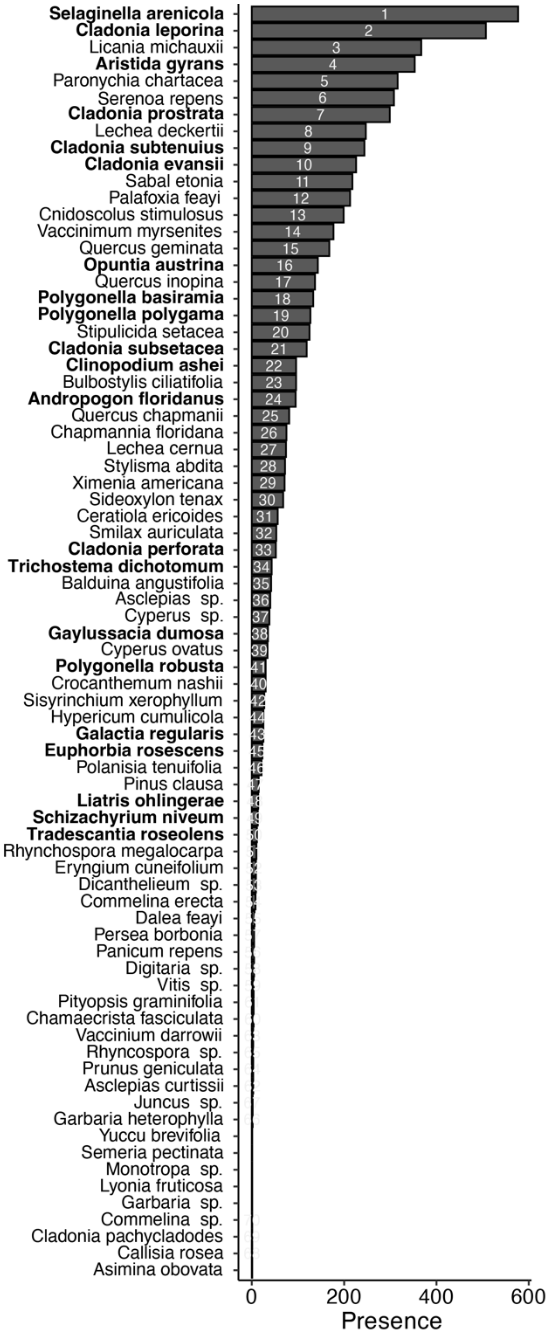


**Supplemental Figure 1.** All plant species surveyed (n = 76) across all study sites (n = 676) organized according to their occurrence rate: presence across all study sites surveyed. Rank order is presented within bars. Species (n = 21) with informative species distribution models (test AUC > 0.7) are in bold.

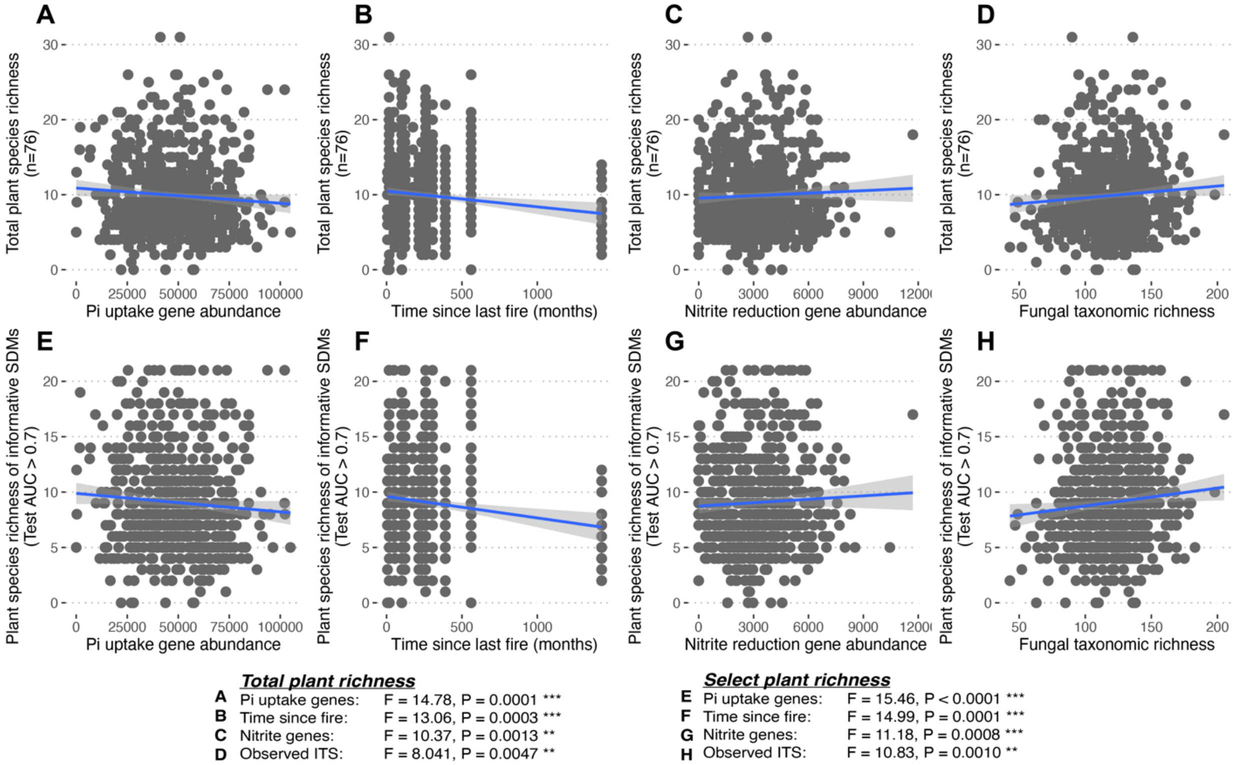


**Supplemental Figure 2.** Significant relationships between abiotic and microbial predictors and plant richness across the landscape. Plant species richness of all identified species in our study (n = 76) as related to **A)** Pi uptake and transport gene (*ugp*) abundance, **B)** time since last fire, **C)** nitrite reducing gene abundance and **D)** observed taxonomic richness of fungi (ITS). Richness of plant species with informative SDMs (n = 21) as related to **E)** Pi uptake and transport gene (*ugp*) abundance, **F)** time since last fire, **G)** nitrite reducing gene abundance and **H)** observed taxonomic richness of fungi (ITS). Blue line represents linear correlation and shaded region represents 95% confidence interval. Statistical significance from generalized linear models generated by stepwise selection is presented for each figure on the bottom.

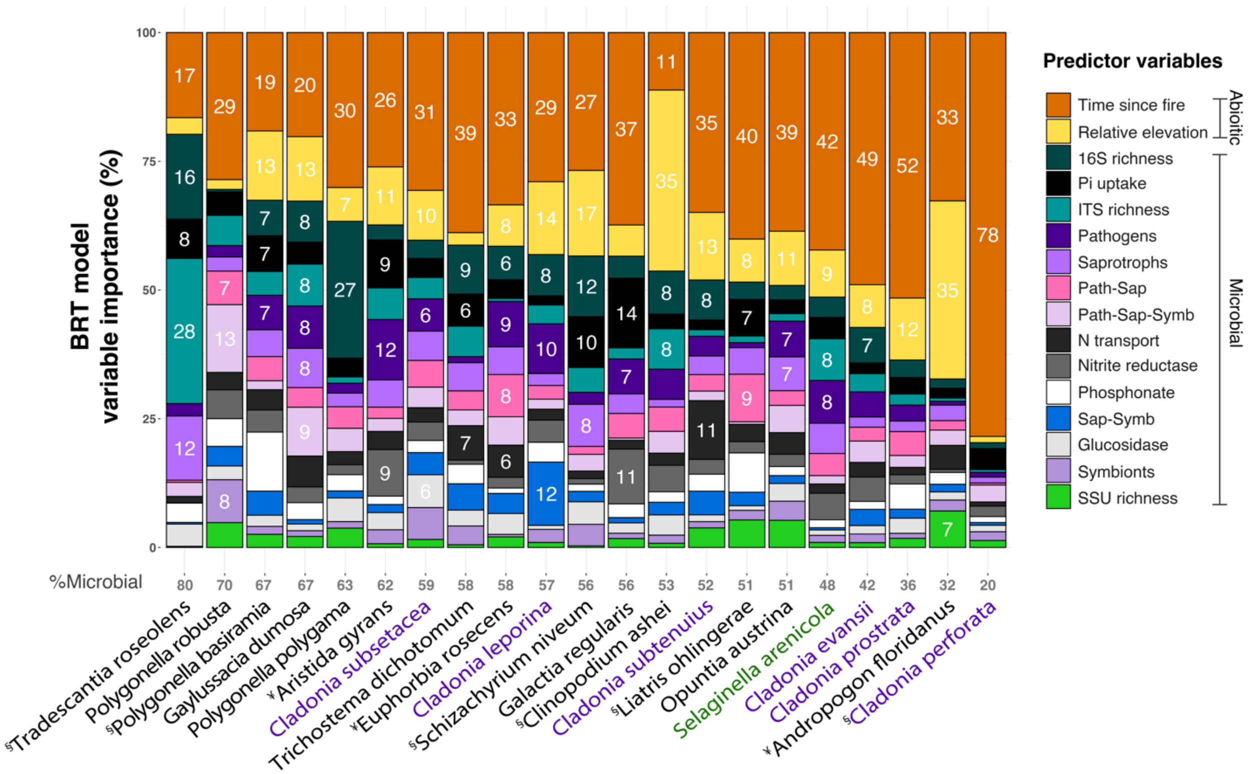


**Supplemental Figure 3.** Total predictor importance for all plant SDMs with test AUC > 0.70, including abiotic variables time since last fire and relative elevation to the water table, as well as all microbial metrics. Total percent of microbial predictor influence on each species' SDM is in gray (bottom). Plant species are arranged along the x-axis in descending order of total relative influence of microbial metrics. Lichens are in purple and the lycopod (*Selaginella arenicola*) is in green. ¥ = plant species endemic to Florida or southeastern USA. § = listed as a threatened or endangered plant species.

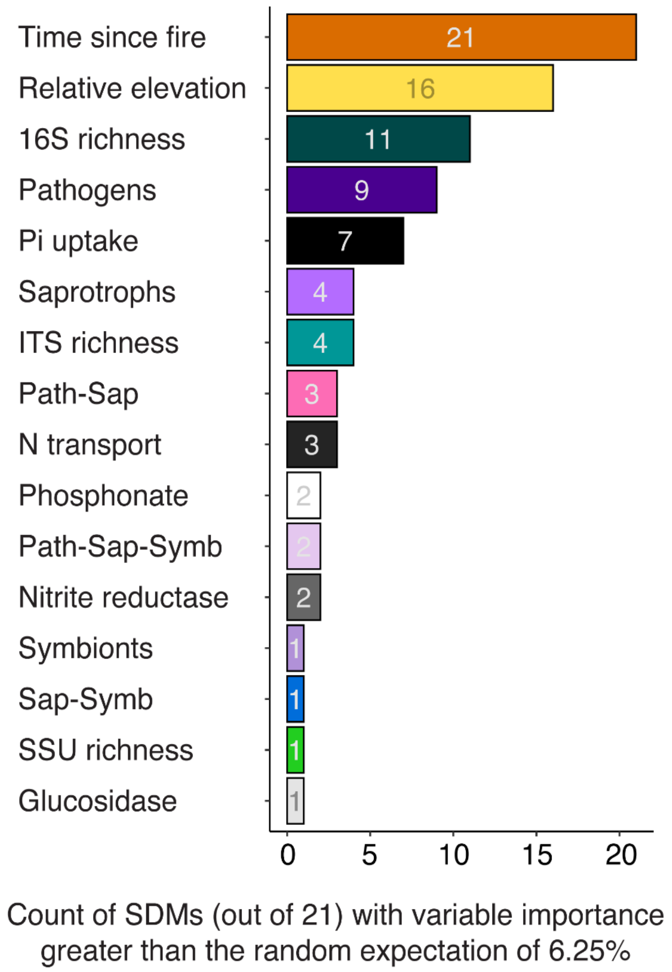


**Supplemental Figure 4.** Summary of predictor variable impact on 21 plant species distribution models (SDMs) with test AUC > 0.70. Count of plant SDMs where the predictor explained greater than the random expectation of 6.25%. For example, time since last fire had a greater impact on 100% of the SDMs than expected by chance, and prokaryotic (16S) species richness had a greater impact on 52% of the SDMs than expected by chance.

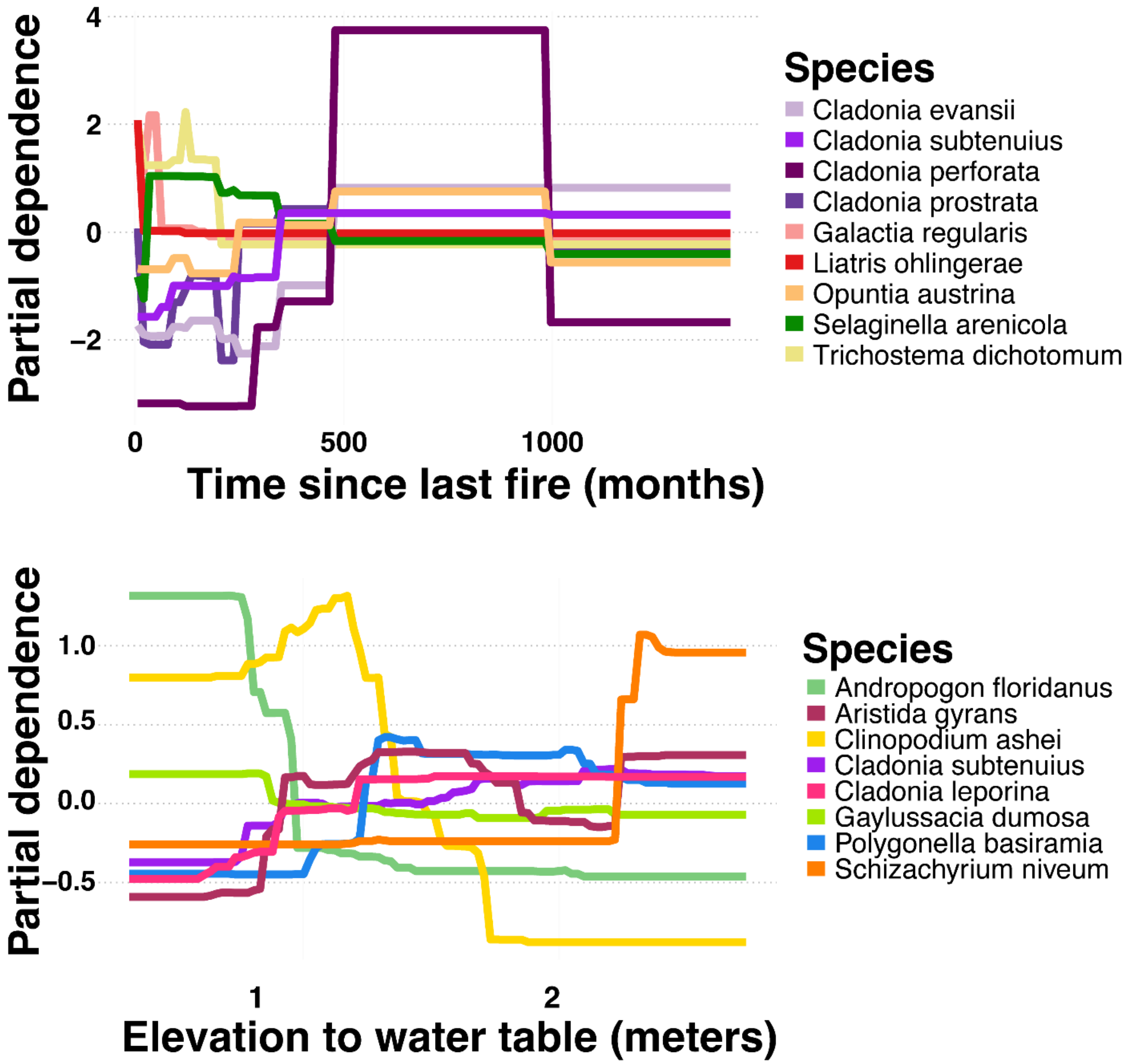


**Supplemental Figure 5.** Partial dependence plots (scaled to a mean of zero to remove effects of overall abundance) for the two abiotic predictors: time since last fire (months) and relative elevation to the water table (meters). Only plant species whose time since last fire variable importance was >35% (mean) and relative elevation variable importance was >11.2% (mean) are shown to emphasize plant taxa with distributions strongly predicted by these abiotic variables. Plant species exhibited a fairly even mix of responses to these abiotic gradients, demonstrating specialization on different parts of these niche axes.

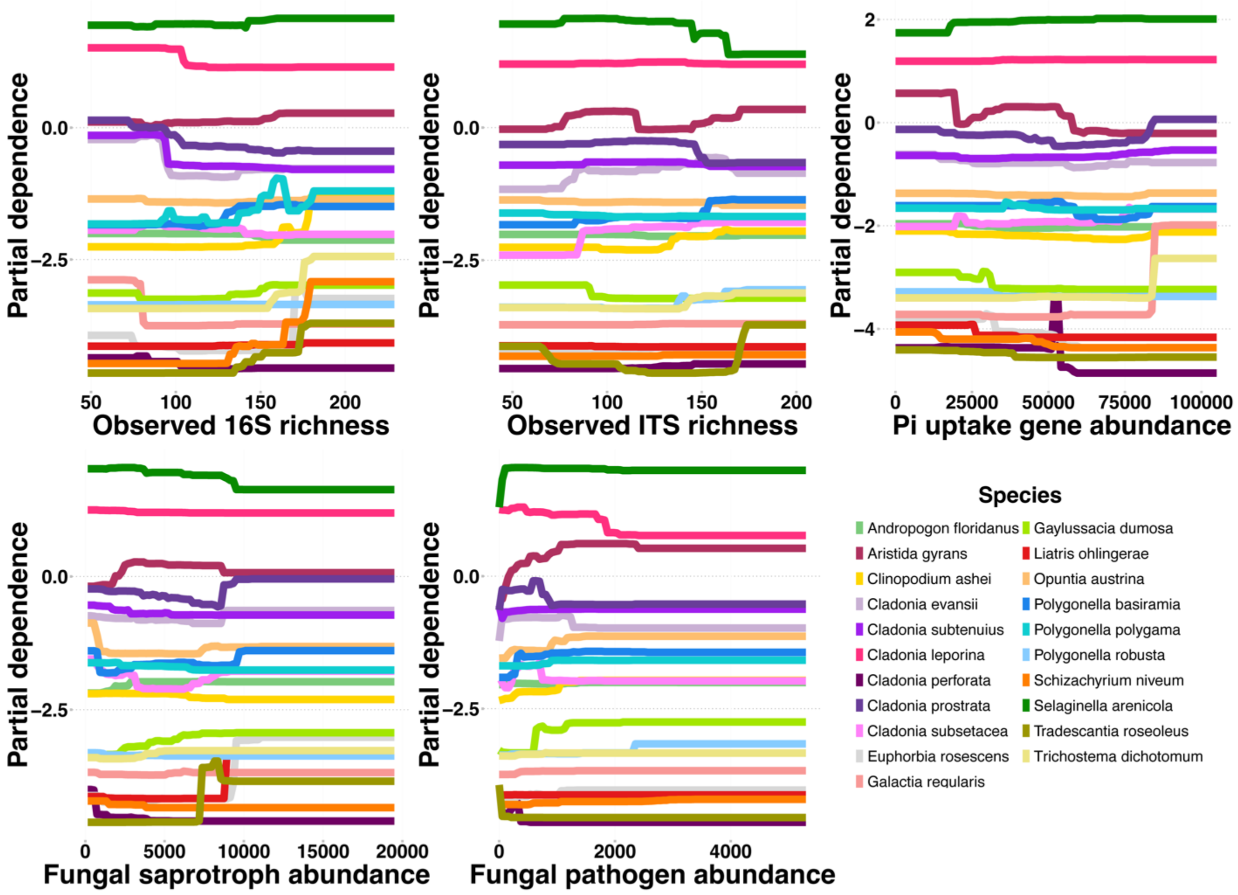


**Supplemental Figure 6.** Partial dependence plots for the five most explanatory microbial predictor variables across the 21 species distribution models with test AUC > 0.70.
